# The neural basis of temperature-driven host seeking in the human threadworm *Strongyloides stercoralis*

**DOI:** 10.1101/2021.06.23.449647

**Authors:** Astra S. Bryant, Felicitas Ruiz, Joon Ha Lee, Elissa A. Hallem

## Abstract

Soil-transmitted parasitic nematodes infect approximately one billion people and are a major cause of morbidity worldwide^1–8^. The infective larvae (iL3s) of these parasites actively search for hosts in a poorly understood, sensory-driven process that requires thermal cues^9,10^. Here, we describe the neural basis of temperature-driven host seeking in parasitic nematodes using the human threadworm *Strongyloides stercoralis*. We show that *S. stercoralis* thermosensation is mediated by the AFD neurons, a thermosensory neuron class that is conserved between parasitic and free-living nematodes^11^. We demonstrate that *S. stercoralis* AFD displays parasite-specific adaptations that enable both nonlinear and linear encoding of temperatures up to human body temperature. Furthermore, we describe a novel thermosensory behavior in which *S. stercoralis* iL3s generate spontaneous reversals of temperature preference at below-body temperatures. Finally, we identify three thermoreceptors selectively expressed in *S. stercoralis* AFD that display parasite-specific sensitivities to human body temperatures and likely enable temperature-driven host seeking by iL3s. Our results are the first direct evidence that the sensory neurons of soil-transmitted parasitic nematodes exhibit parasite-specific neural adaptations and sensory coding strategies that allow them to target human hosts, a finding with important implications for efforts to develop new therapeutic strategies for nematode control.

---

Soil-transmitted gastrointestinal helminths such as the human threadworm *Strongyloides stercoralis* and the hookworms *Ancylostoma duodenale* and *Necator americanus* are a major source of neglected disease affecting the world’s most socioeconomically depressed communities^1–8^. These parasites can cause chronic gastrointestinal distress as well as stunted growth and cognitive impairment in children. *S. stercoralis* is estimated to infect at least 610 million people worldwide^12^; infections can be fatal for immunosuppressed individuals^13–15^. Current therapeutics against parasitic nematodes exclusively target ongoing infections. These anthelmintic drugs have variable efficacy at clearing infections, and the lack of prophylactic treatments results in high reinfection rates^16–19^. Additionally, repeated dosage with anthelmintics may give rise to drug-resistant nematodes^16–18^, underscoring the need for preventative strategies. A mechanistic understanding of how parasitic nematodes locate hosts may enable the development of new therapeutic approaches to preventing infections.

For parasitic nematodes that infect humans and other mammals, host detection and subsequent infection require thermal cues^10,20^. Mammalian-parasitic iL3s, including those of *S. stercoralis*, display robust attraction to mammalian body heat when exposed to temperatures above an ambient cultivation temperature (T_C_; Fig. 1a)^10,11^. This positive thermotaxis behavior is specific to mammalian-parasitic nematodes: insect parasitic nematodes and the free-living model nematode *Caenorhabditis elegans* are not attracted to mammalian body heat^10^. Indeed, exposure to temperatures above T_C_ drives either negative thermotaxis or noxious heat avoidance by *C. elegans* adults^21–26^. The molecular and neural basis of *C. elegans* thermosensation has been extensively studied^21,24,26,27^. However, virtually nothing is known about the neural mechanisms that underlie thermosensory behaviors in parasitic nematodes or any other endoparasitic animal. More generally, although sensory neuroanatomy is broadly conserved across free-living and parasitic nematode species^9^, the neural mechanisms that enable nematodes to engage in species-specific behaviors such as host seeking have not been identified.

**Figure 1.**
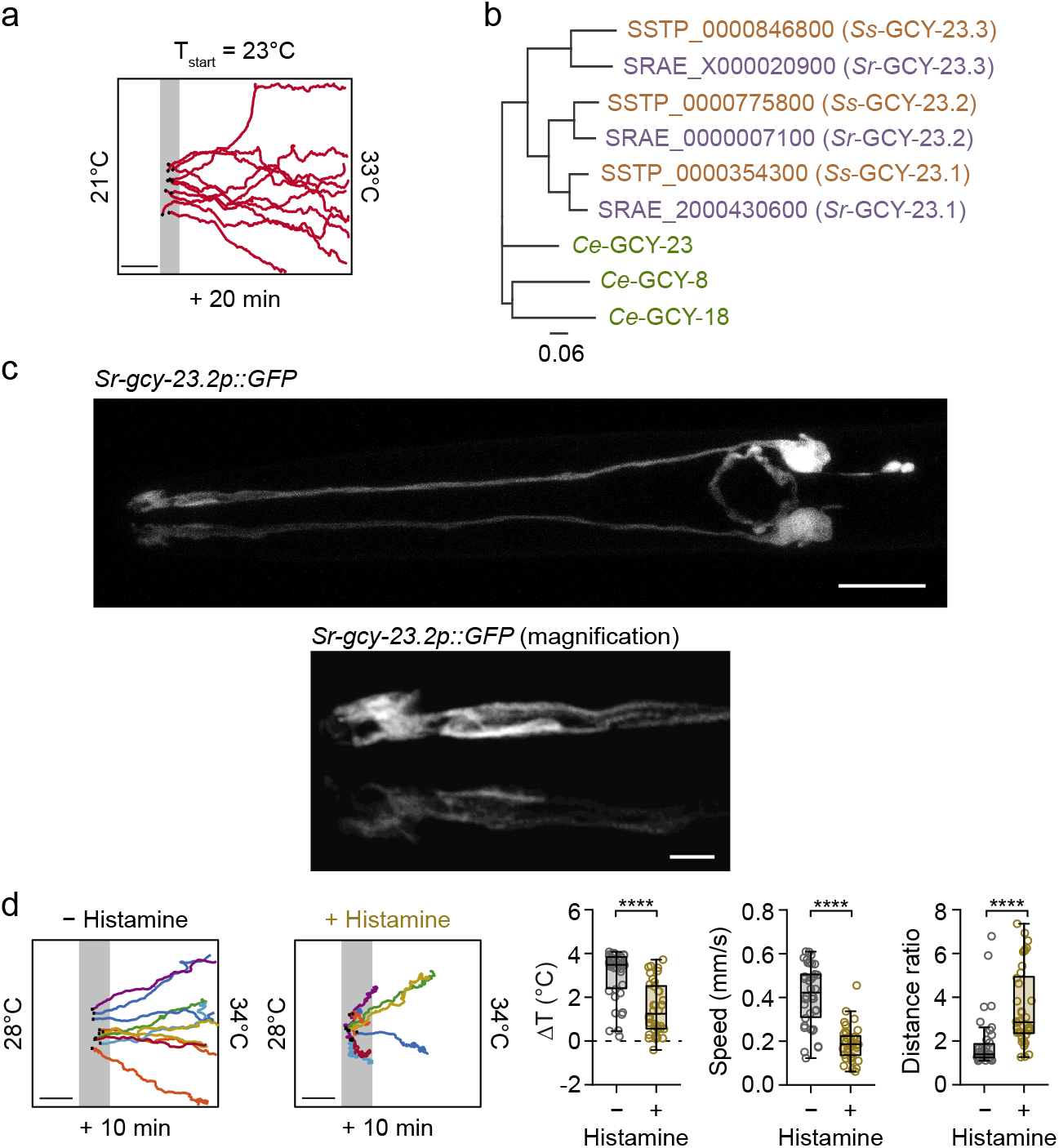
The *S. stercoralis* AFD neurons are required for positive thermotaxis toward host body temperatures. **a.** Tracks of *S. stercoralis* iL3s engaging in positive thermotaxis toward host body temperatures. Worms were placed at 23°C in a ∼21-33°C gradient and allowed to migrate for 20 min. **b.** Phylogenetic tree of *C. elegans* AFD-specific receptor-type guanylyl cyclases (AFD-rGCs, green text) and homologous *S. stercoralis* (orange text) and *S. ratti* (purple text) proteins. Putative AFD-rGCs for *Strongyloides spp*. were identified based on protein sequence homology with *C. elegans* GCY-8, GCY-18, and GCY-23. SSTP and SRAE numbers indicate *S. stercoralis* and *S. ratti* WormBase ParaSite gene IDs, respectively; gene names indicating homology to *C. elegans* GCY-23 (GCY-23.1, GCY-23.2, and GCY-23.3) for *S. stercoralis* and *S. ratti* sequences are shown in parentheses. **c.** The promoter region for *Sr-gcy-23.2* drives GFP expression in a single neuron pair (top) with a dendritic morphology (bottom left) that matches the lamellar morphology of *S. stercoralis* ALD^11,32,33^. For top image, scale bar is 10 m. For bottom left image, scale bar is 2 m. For all images, anterior is to the left. **d.** HisCl1-mediated silencing of the *S. stercoralis* AFD neuron pair results in profound reductions in positive thermotaxis by iL3s. *Left:* Tracks of *Sr-gcy-23.2p::strHisCl1*; *Sr-gcy-23.2p::GFP* iL3s treated with BU saline or histamine (either 20 or 50 μL) in BU for at least 1 h, migrating in a ∼22-34°C gradient for 10 min. T_start_ = 30°C. *Right*: Change in temperature, average speed, and distance ratio (total distance ÷ maximum displacement) exhibited by individual iL3s. n = 38 worms for iL3s treated with BU saline, 36 worms for histamine-treated iL3s; \*\*\*\**p*<0.0001, two-sided Mann-Whitney test. Exact *p* values for all figures are provided in Supplemental Data File 1. For all plots of worm tracks: Scale bar = 2 cm, gray zone = 1°C centered at T_start_. A subset of 10 randomly selected tracks are plotted and only a portion of the full gradient is shown.

## Genetic identification of *S. stercoralis* thermosensory neurons

To understand the neural mechanisms of thermosensation in *S. stercoralis* iL3s, we first used a genetic approach to identify the primary thermosensory neurons of *S. stercoralis*. The identification of thermosensory neurons in *S. stercoralis* has been an historical challenge due to morphological differences in sensory neuroanatomy that complicate anatomical comparisons between *S. stercoralis* and *C. elegans*^11^. In *C. elegans*, the AFD sensory neuron pair provides the primary thermosensory drive for thermotaxis navigation^26,28–31^. *C. elegans* AFD neurons are structurally defined by “finger-like” microvilli that emerge from the dendritic ends at the nose of the animal and are thought to enhance neuronal sensitivity to temperature fluctuations^26^. *S. stercoralis* lacks sensory neurons with these “finger-like” structures^32^. The closest anatomical homologs appear to be the multi-layered “lamellae” of the ALD neurons, which are the only elaborate sensory endings in the *S. stercoralis* amphid sensory organs^32^. However, due to both variations in the arrangement of the cell bodies that comprise the amphid sensory organs (13 soma for *S. stercoralis* versus 12 soma for *C. elegans*) and life stage-specific changes in the morphology of sensory neurons, whether ALD neurons are homologs of the *C. elegans* AFD thermosensory neurons or AWC chemosensory neurons (the other *C. elegans* amphid neurons with elaborate sensory endings) has been unclear^11,32–34^.

In *C. elegans*, the receptor-type guanylyl cyclases (rGCs) GCY-8, GCY-18, and GCY-23 are expressed selectively in *C. elegans* AFD, contribute to thermosensation via activation of the cyclic GMP-gated channel subunit TAX-4, and are used as AFD-specific genetic markers^31,35–38^. The *S. stercoralis* TAX-4 homolog, *Ss-*TAX-4, is required for positive thermotaxis by iL3s^10,20^. Thus, parasite homologs of the AFD-rGCs may serve as molecular markers for AFD-specific function, permitting the genetic identification of the sensory neurons that drive the temperature-driven movements of *S. stercoralis* iL3s. The use of parasite homologs is necessary as *C. elegans* transgenes and regulatory elements are generally inactive in *Strongyloides* species^39–41^.

We used BLASTP to identify homologs of the three *C. elegans* AFD-rGCs in *S. stercoralis* and the closely related rat parasite *Strongyloides ratti*. The genomes of *S. stercoralis* and *S. ratti* each contain three putative AFD-rGCs (Fig. 1b). A phylogenetic comparison between the nine protein sequences revealed that although there is clear 1-to-1 homology between the *Strongyloides* species, all six *Strongyloides* sequences are most homologous to *C. elegans* GCY-23 (Fig. 1b, Fig. S1a). Based on protein sequence homology, we therefore named the *S. stercoralis* and *S. ratti* homologs GCY-23.1, GCY-23.2, and GCY-23.3. We then extracted promoter sequences for the three *S. stercoralis* rGCs, as well as *Sr*-GCY-23.2, and found that all four promoters drive strong transgene expression in a single *Ss-tax-4*-positive head neuron pair in *S. stercoralis* iL3s (Fig. 1c, Fig. S1b,c). Using confocal microscopy, we found that this neuron pair displays elaborate structures at the nose tip that strongly resemble electron microscopy reconstructions of the ALD lamellae (Fig. 1c)^33,34^. Based on genetic homology, we thus identified the *S. stercoralis* ALD neurons as homologs of the *C. elegans* AFD neurons (and hereafter refer to them as *S. stercoralis* AFD neurons, or *Ss-*AFD).

## *S. stercoralis* AFD neurons are required for heat seeking

To test whether the *Ss*-AFD neurons are required for the attraction of *S. stercoralis* iL3s to mammalian body temperatures, we chemogenetically silenced them via AFD-specific expression of the *Drosophila* histamine-gated chloride channel HisCl1^42^, which we codon-optimized for *Strongyloides* (*Sr-gcy-23.2p::strHisCl1*, Fig. S2a). We found that silencing *Ss*-AFD reduced the migration of iL3s toward temperatures approximating mammalian body heat, blocked temperature-driven increases in worm speed, and reduced thermal drive-dependent inhibition of local search (Fig. 1d, Fig. S2b). Our results establish single neuron genetic targeting for the first time in a non*-Caenorhabditis* nematode and demonstrate that the *Ss*-AFD neurons are required for the heat-seeking behavior of iL3s, thus confirming the impact of *Ss*-ALD laser ablation^33^. Notably, our results indicate that *S. stercoralis* and *C. elegans* rely on a homologous primary sensory neuron pair to drive distinct behaviors in response to temperatures above ambient: negative thermotaxis by *C. elegans* adults^26^ and positive thermotaxis by *S. stercoralis* iL3s.

## Novel encoding of naturalistic warming stimuli by *S. stercoralis* AFD neurons

Do temperature-driven responses in *Ss*-AFD display species-specific adaptations, or is AFD encoding of temperature conserved between parasitic and free-living nematodes? We generated iL3s expressing yellow cameleon YC3.60^43^ in *Ss*-AFD, which allowed us to image neural activity for the first time in a non-*Caenorhabditis* nematode. We designed thermal stimuli that mimic the temperatures experienced by iL3s engaged in positive thermotaxis toward host body temperatures; these naturalistic warming stimuli were delivered to immobilized worms via a custom PID-controlled thermal stimulator (0.025 °C/s; Fig. S3a,b). We then imaged from *S. stercoralis* AFD and *C. elegans* AFD under the same experimental conditions. Consistent with previous findings^26,36,44,45^, naturalistic warming stimuli drive large depolarizations in *C. elegans* AFD (*Ce*-AFD) above a response threshold (T*_AFD_) set near cultivation temperature (T_C_; Fig. 2a,b). *Ce*-AFD monotonically and nonlinearly encodes temperature deviations near T_C_, such that the lowest *Ce*-AFD response is driven by the coolest temperature in the thermal stimulus (Fig. 2c) and the *Ce*-AFD response is positively correlated with increasing temperature only in a narrow temperature range near T*_AFD_ and not at warmer temperatures (Fig. 2d-h).

**Figure 2.**
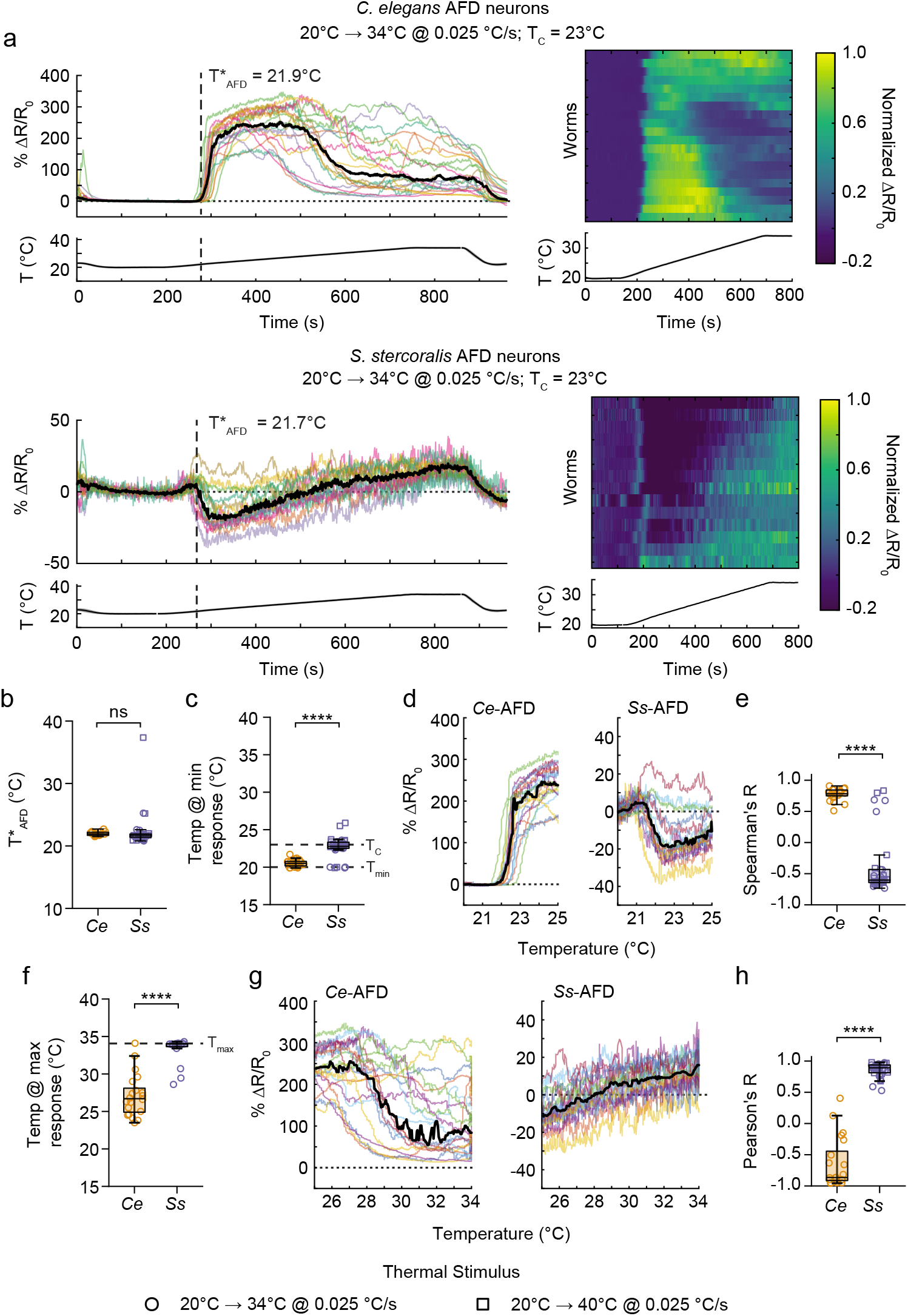
Naturalistic warming ramps drive parasite-specific warming-triggered hyperpolarizations followed by near-linear thermal encoding in *S. stercoralis* AFD neurons. **a.** *C. elegans* AFD and *S. stercoralis* AFD responses to a 20-34°C warming temperature ramp that approximates the temperatures experienced by iL3s engaging in positive thermotaxis in a ∼22-34°C gradient. Colored traces show the responses of individual worms; black traces show median responses. Heatmaps show responses normalized to the maximum ΔR/R_0_ response for each species, with R_0_ set to zero. Responses are ordered by hierarchical cluster analysis. For all plots, temperature traces show the average recorded temperature (mean in solid line, SD in shading; in some cases, shading is too small to be visible). Warming ramp slope = 0.025 °C/s. R_0_ (horizontal dotted lines) = response at 20°C. n = 15 worms for *S. stercoralis*, 20 for *C. elegans*. **b.** Thermal threshold (T*_AFD_) of *C. elegans* and *S. stercoralis* AFD calcium responses, where T*_AFD_ is defined as the first temperature where R/R_0_ is significantly different from baseline (3 * STD R_0_) for at least 10 seconds. Data are responses to either a 20-34°C (circles) or a 20-40°C (squares) warming ramp. Meaningful differences between *C. elegans* T*_AFD_ and *S. stercoralis* T*_AFD_ were not detected (difference between median T*_AFD_ = −0.2°C (*Ss-*AFD vs *Ce*-AFD), effect size = 0.187, power = 0.095, Mann-Whitney test). **c.** Temperature that elicits the lowest calcium response in *Ce*-AFD and *Ss*-AFD, in response to either a 20-34°C (circles) or a 20-40°C (squares) warming ramp. Horizontal dashed lines indicate T_C_ (23°C) and T_min_ (20°C). \*\*\*\**p*<0.0001, Mann-Whitney test. **d.** Plots of % ΔR/R_0_ as a function of temperature at near-ambient temperatures (20-25°C). Horizontal dotted line indicates R_0_ (response at 20°C). **e.** Quantification of monotonic correlation between temperature and calcium responses in the 20-25°C range. Values are Spearman’s correlation coefficients. \*\*\*\**p*<0.0001, Mann-Whitney test. **f.** Temperature that elicits the highest calcium response in *Ce*-AFD and *Ss*-AFD, in response to a 20-34°C warming ramp. Horizontal dashed line indicates the maximum temperature tested (T_max_; 34°C). \*\*\*\**p*<0.0001, Mann-Whitney test. n = 15 worms for *S. stercoralis*, 20 for *C. elegans*. **g.** Plots of % ΔR/R_0_ as a function of temperature at above-ambient temperatures (25-34°C). Horizontal dotted line indicates R_0_ (response at 20°C). **h.** Quantification of linear correlation between temperature and calcium responses at 25°C and higher. Values are Pearson’s correlation coefficients. \*\*\*\**p*<0.0001, Mann-Whitney test. For all plots: icons indicate responses of individual worms, boxes show median and interquartile ranges, whiskers show min and max values. T_C_ (ambient) = 23°C. n = 20 worms for *C. elegans*, 30 for *S. stercoralis*, unless otherwise specified. All Mann-Whitney tests are two-sided.

In contrast, temperature-driven responses in *Ss*-AFD encode different features of the thermal stimulus and display multiple parasite-specific adaptations. *Ss*-AFD also has a response threshold near T_C_ (Fig. 2a,b). However, the *Ss*-AFD response at T*_AFD_ is dominated by a warming-triggered inhibition, such that the temperature that elicits the lowest *Ss*-AFD response is near T_C_ (Fig. 2c). This warming-triggered inhibition results in a negative monotonic relationship between temperature and *Ss*-AFD activity near T_C_ (Fig. 2d,e). Following warming-triggered inhibition, *Ss*-AFD responses show near-linear positive encoding of temperatures ranging from near T_C_ to above human body temperature, with peak excitatory responses driven by the warmest temperature in the stimulus (Fig. 2f-h, Fig. S4). These parasite-specific response properties provide a mechanism for *Ss*-AFD to encode additional information and differentiate between a broad range of host and environmental temperatures, and are likely critical for the ability of iL3s to host seek over relatively long distances^9^. The observation that natural behaviors arise from species-specific functional adaptations in developmentally homologous neural structures is a central tenet of neuroethology. However, the nature of such functional adaptations remains poorly understood. Our results demonstrate that the parasitic behaviors of soil-transmitted nematodes emerge from functional adaptations at the earliest stages of sensory information processing within the conserved nematode nervous system.

Next, we tested the response of *Ss*-AFD and *Ce*-AFD to rapid warming temperature ramps (0.1 °C/s; Fig. S5a-c). Consistent with previous reports^46^, *Ce*-AFD responses are similar across ramp rates. In contrast, the shape of *Ss*-AFD responses is partially dependent on ramp steepness. Rapid temperature ramps are less likely to produce warming-triggered inhibition, such that for the majority of iL3s, the coolest temperature in the rapid stimulus ramp drives the lowest *Ss*-AFD response (Fig. S5c-e). T*_AFD_ is not affected by ramp speed (Fig. S5f). Furthermore, *Ss*-AFD maintains both near-linear positive encoding of temperature above ∼T_C_ in response to rapid ramp speeds and a peak Ca^2+^ response at the warmest temperatures in the stimulus (Fig. S5g,h). These results suggest that the mechanisms of warming-triggered inhibition in *Ss*-AFD may be partially separate from the mechanisms of warming-triggered depolarizations and the setting of thermal response threshold. Warming-triggered inhibitory responses have not been reported previously in nematodes and thus may represent a parasite-specific molecular mechanism for encoding thermosensory signals. Taken together, our observations highlight the potential of elements in the *Ss*-AFD thermotransduction pathway as novel targets for parasite-specific therapeutic interventions.

## *Ss*-AFD responses are regulated by cultivation temperature

The thermotaxis behaviors of both *C. elegans* and *S. stercoralis* are regulated by changes in cultivation temperature^11,26^. In *C. elegans*, exposing animals to a new cultivation temperature shifts the thermal threshold of AFD toward the new T_C_^26,36,44,45^. To determine whether responses in *Ss*-AFD and *Ce*-AFD display similar T_C_-dependent changes, we cultivated worms overnight at 15°C, then exposed them to naturalistic thermal stimuli that were shifted toward cooler temperatures (Fig. S3c). Cultivation at 15°C shifts T*_AFD_ toward cooler temperatures in both *Ss*-AFD and *Ce*-AFD (Fig. S6a-d). The downward shift in thermal threshold does not eliminate the unique properties of the *Ss*-AFD response. The response at the cooler T*_AFD_ is still dominated by warming triggered inhibition, with a peak hyperpolarization just above T*_AFD_ (Fig. S6e), and a peak depolarization in response to the warmest temperature in the stimulus (Fig. S6f). Notably, despite the shift in T*_AFD_ toward cooler temperatures, *Ss*-AFD still responds near-linearly to temperatures up to host body temperatures (Fig. S6c), consistent with earlier behavioral results demonstrating that *S. stercoralis* iL3s cultivated at 15°C engage in robust positive thermotaxis toward skin temperatures^10^. The preservation of near-linear encoding at host body temperatures, combined with experience-dependent expansions in the dynamic range of *Ss*-AFD thermosensitivity, further supports a model in which parasite-specific adaptations in sensory information processing pathways actuate the ability of iL3s to host seek across a range of ethological contexts.

## Characterizing negative thermotaxis behaviors of *S. stercoralis* iL3s

Besides attraction to host body temperatures, mammalian-parasitic iL3s display negative thermotaxis: migration toward cooler temperatures when exposed to temperatures below their ambient cultivation temperature^10^. Previously, negative thermotaxis was measured exclusively using iL3s exposed to a ∼20-34°C gradient^10^; here we expand on those initial observations by measuring migration in a ∼12-22°C gradient (Fig. S7). When placed at 20°C or 17°C, iL3s cultivated at 23°C display negative thermotaxis, accumulating near ∼16°C (Fig. S7a). These migration patterns are regulated by recent experience, such that iL3s cultivated at 15°C for at least 2 hours accumulate at cooler temperatures than iL3s cultivated at 23°C (Fig. S7b-d). Does accumulation near ∼16°C represent a *bona fide* preferred temperature? If yes, iL3s placed below 16°C should navigate up thermal gradients toward their “preferred” temperature. In fact, iL3s placed at 14°C in a ∼12-22°C gradient remain at cooler temperatures, regardless of cultivation temperature (Fig. S7). Thus, *S. stercoralis* iL3s do not appear to have a single preferred “cool” temperature. Rather, negative thermotaxis appears to be regulated by the strength of thermal drive, which decays as a function of deviations from T_C_, similar to positive thermotaxis^10^.

## *Ss*-AFD mediates negative thermotaxis via an *Ss-tax-4* signaling cascade

Positive and negative thermotaxis behaviors represent distinct iL3 behavioral modes, and it was not known whether these behaviors are regulated by the same signaling cascades and primary sensory neurons. We found that CRISPR/Cas9-mediated targeted mutagenesis of *Ss-tax-4*^47^ and HisCl1-mediated silencing of *Ss*-AFD activity both abolish negative thermotaxis by iL3s (Fig. S8a,b), similar to positive thermotaxis (Fig. 1d)^10^. In *C. elegans*, positive and negative thermotaxis behaviors toward T_C_ also rely on *Ce-tax-4*-dependent sensory transduction in *Ce*-AFD^26,36,48,49^. Thus, our results provide additional evidence that the opposing thermosensory behaviors of *S. stercoralis* iL3s and *C. elegans* adults reflect adaptations of a broadly conserved nematode thermosensory circuit.

Next, we tested the effect of cooling temperature ramps on *Ss*-AFD activity, compared to *Ce*-AFD. Delivery of a rapidly cooling temperature ramp (range: 22-13°C, Fig. S8c), drives near-linear cooling-mediated hyperpolarizations in both *Ce*-AFD and *Ss*-AFD (Fig. S8d,e). These results suggest that cooling-mediated inhibition of AFD corresponds to opposing behaviors in free-living and parasitic nematodes: negative thermotaxis toward below-ambient temperatures for *S. stercoralis* iL3s versus positive thermotaxis toward the “remembered” cultivation temperature for *C. elegans* adults.

## *S. stercoralis* iL3s display a unique “reversal” behavior

When navigating in the environment, *S. stercoralis* iL3s likely encounter temperature gradients that terminate well below host body temperature and do not reflect the presence of a host animal (*e.g*., sun-warmed soil). Do iL3s have a mechanism that enables them to avoid accumulation at these “false positive” heat sources? To address this question, we examined the behavior of iL3s in temperature gradients that are warmer than ambient but well below host body heat. We first exposed iL3s cultivated at 23°C to temperature ramps ranging from 15-25°C (T_start_ = 23°C). Under these conditions, most iL3s initially engage in positive thermotaxis and accumulate at the warmest temperature in the gradient, consistent with the behaviors of iL3s in thermal gradients that end near host body temperature^10^. However, unlike our observations with warmer gradients, we found that some of these heat-seeking iL3s will, within minutes, abruptly transition to sustained negative thermotaxis (Fig. 3a; median percent of heat seeking iL3s that reverse = 35%; median dwell time at T_max_ = 216 s). Worms cultivated at 15°C display similar “reversal” behaviors (Fig. 3b; median dwell time at T_max_ = 126 s), with increased reversal frequencies observed in thermal gradients that terminate closer to T_C_ (Fig. 3c; median percent of heat-seeking iL3s that reverse in a 15-25°C gradient = 3.46%, median percent of heat-seeking iL3s that reverse in a 12-22°C gradient = 97.25%). Across cultivation temperatures, reversal behaviors were only ever observed in thermal gradients that end well below host body temperature^10^. Thus, the likelihood that iL3s will transition from positive to negative thermotaxis is modulated by both cultivation temperature and the assay gradient (Fig. 3c). This rapid reversal of thermal preferences appears to be parasite-specific, as *C. elegans* behavioral preferences only change on the timescale of hours, not minutes^21,26,50^. Reversal behaviors likely allow iL3s to rapidly abandon an erroneous attraction to non-host heat sources. Our results identify a new component of temperature-driven host seeking, and indicate that soil-transmitted parasitic nematodes rely on a host-seeking strategy that flexibly adjusts in response to sensory experience.

**Figure 3.**
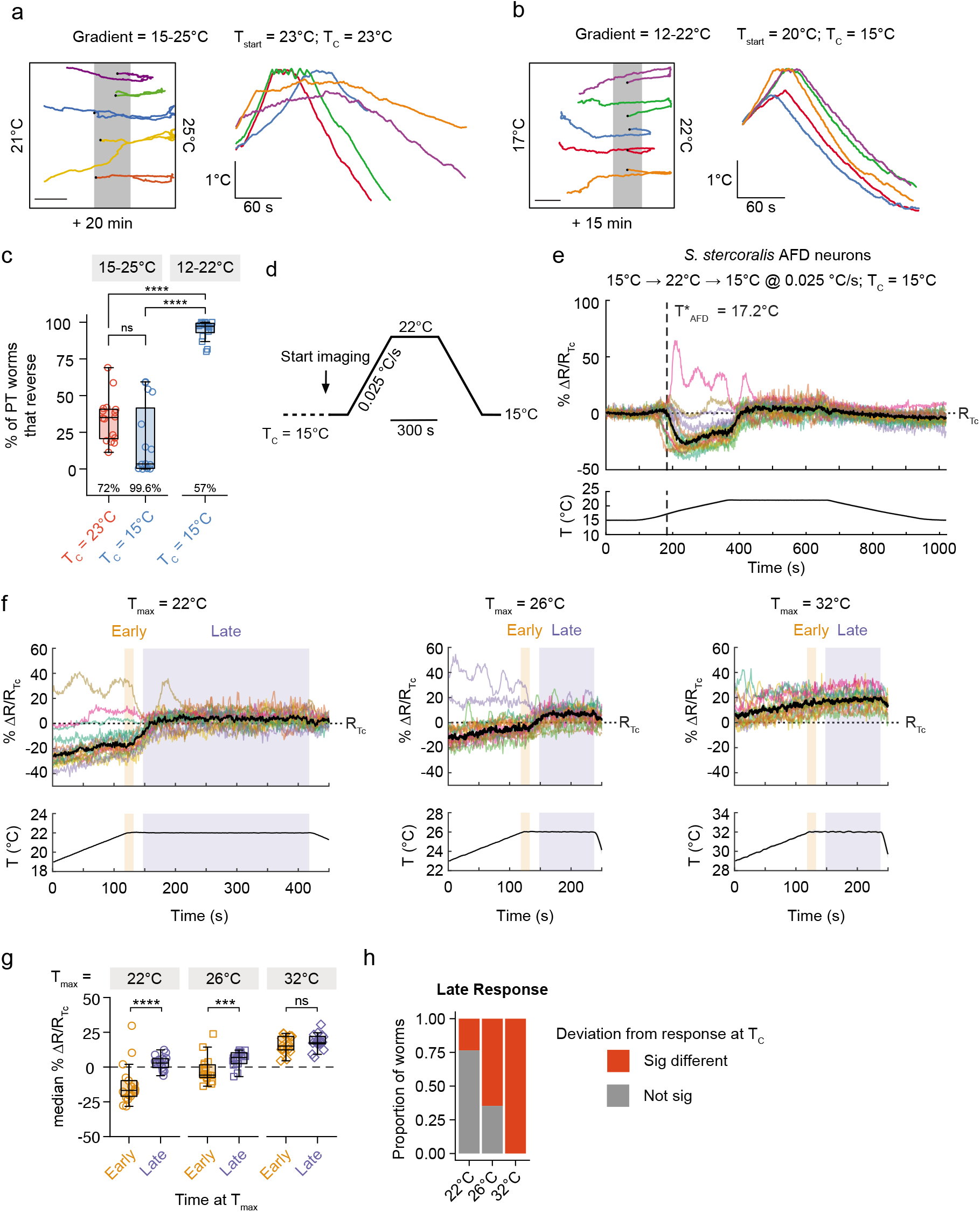
*S. stercoralis* iL3s display rapid behavioral and neural adaptation in response to thermosensory cues well below host body temperatures. **a.** Example tracks of individual *S. stercoralis* iL3s engaging in “reversal behavior,” which is characterized by an abrupt transition from sustained positive thermotaxis to sustained negative thermotaxis. Gradient range: 15-25°C. T_start_ = 23°C. T_C_ = 23°C. Left: Example worm tracks artificially distributed along the y-axis to increase visual clarity. Right: Temperatures experienced by individual iL3s as a function of time; traces are the same worms displayed in the left panel. Median time spent near T_max_ = 216 s (IQR = 166-244 s). n = 5 worms. **b.** Worms cultivated at 15°C also display reversal behaviors. Gradient range: 12-22°C. T_start_ = 20°C. T_C_ = 15°C. Median time spent near T_max_ = 126 s (IQR = 106-150 s). n = 9 worms, plots show subset of tracks. Plots use same conventions as in panel **a**. **c.** Percent of *S. stercoralis* iL3s engaging in positive thermotaxis (PT) that reverse into negative thermotaxis. Annotations along the x-axis are the average percentage of the total population that engages in positive thermotaxis under each experimental condition. Boxes indicate medians and interquartile ranges; whiskers indicate min and max values. n = 15 assays for the 15-25°C gradient when T_C_ = 15°C, 16 for all other conditions. ns = not significant (*p*>0.05), \*\*\*\**p*<0.0001, Kruskal-Wallis test with Dunn’s post-test. **d.** Diagram of thermal stimulus that mimics temperature ramps experienced by 15°C-cultivated iL3s displaying reversal behavior in a 12-22°C gradient. These parameters were chosen because they are the assay conditions most likely to elicit reversal behaviors. Ramp slope: ±0.025 °C/s. Holding time at 22°C = 5 min. **e.** *Ss-*AFD calcium responses to the thermal stimulus shown in panel **d**. For calcium responses, colored lines indicate responses from individual worms, black line indicates median response. Temperature traces are mean + SD recorded temperature (lines + shading; in some cases, shading is too small to be visible). Dashed vertical line indicates thermal threshold. Dotted horizontal line indicates baseline temperature response, equivalent to the response at T_C_ (R_Tc_). n = 17 worms. T_C_ = 15°C. **f.** *Ss-*AFD calcium responses to different maximum temperatures (T_max_): 22°C, 26°C, and 32°C. Colored shading indicates time windows used for analyses in panels **g**, **h**. n = 17 worms (22°C or 26°C), 16 worms (32°C). T_C_ = 15°C. All other conventions are the same as panel **e**. **g.** Adaptation of the steady-state *Ss*-AFD calcium responses at different T_max_ values. Calcium responses are calculated over two analysis windows: the early T_max_ response (first 15 seconds) and the late T_max_ response (30 seconds - end). ns = not significant (*p*>0.05), \*\*\**p*<0.001, \*\*\*\**p*<0.0001, two-way repeated-measures ANOVA with Šidák’s multiple comparisons tests. Boxes indicate medians and interquartile ranges; whiskers indicate min and max values. n = 17 worms (22°C or 26°C), 16 worms (32°C). T_C_ = 15°C. **h.** Temperature stimuli that are more likely to result in reversal behaviors (*i.e*., T_max_ = 22°C and T_max_ = 26°C vs T_max_ = 32°C) result in a higher proportion of late T_max_ responses that are not significantly different from responses to T_C_. For each worm, significant T_max_ responses are those that vary from the average T_C_ response by at least 3 standard deviations. 22°C vs 26°C: *p*>0.05; 26°C vs 32°C: *p*>0.05; 22°C vs 32°C: *p*<0.0001, Fisher’s exact tests with Bonferroni-Dunn correction for multiple comparisons. T_C_ = 15°C. All statistical tests are two-sided; when possible we report multiplicity adjusted *p* values.

To test for neural correlates of the reversal behavior, we designed imaging stimuli that mimicked reversal trajectories (Fig. 3d), then quantified *Ss*-AFD responses at the transition between an initial warming ramp and sustained exposure to a maximum temperature (T_max_). We found that *Ss*-AFD responses at T_max_ rapidly (within ∼30 s) adapted toward the baseline T_C_ response (Fig. 3e-h; median early response at T_max_ = −16.8% ΔR/R_Tc_, median late response T_max_ = 2.9% R/R_Tc_). To test whether this rapid adaptation may be a neural correlate of reversal behavior, we compared *Ss*-AFD responses to a range of T_max_ values associated with a range of reversal rates: 22°C (many reversals), 26°C (some reversals) and 32°C (no reversals). We found that the degree of *Ss*-AFD response adaptation approximates the likelihood that thermal stimuli would drive reversal behaviors: for iL3s cultivated at 15°C, *Ss*-AFD responses to steady-state thermal stimuli strongly adapt at 22°C, moderately adapt at 26°C, and do not adapt at 32°C (Fig. 3f-h). This result suggests that rapid adaptation in the *Ss*-AFD response to some, but not all, temperature stimuli may underlie the ability of iL3s to selectively engage in reversal behaviors.

Most of the highly adapting *Ss*-AFD responses are warming-triggered hyperpolarizations. We therefore tested whether rapid adaptation reflects a general inability of *Ss*-AFD to maintain prolonged hyperpolarization. Cooling temperature ramps from 22-13°C drive *Ss*-AFD hyperpolarizations of similar magnitude to non-adapted warming-triggered hyperpolarizations at 22°C, but do not display rapid adaptation (Fig. S9a,b). These results suggest that *Ss*-AFD response adaptation is likely a consequence of the thermal stimuli themselves rather than the direction of the *Ss*-AFD response, and that distinct processes cause cooling- and warming-triggered hyperpolarizations in *Ss*-AFD. In *C. elegans*, although *Ce*-AFD can display minutes-long adaptation to recent temperature experiences^26,36,46,51^, these changes do not appear to affect behavioral temperature preferences, which are thought to adapt primarily on longer (hours-long) timescales^26,46^. Thus, our results suggest that iL3 reversal behavior may reflect a parasite-specific behavioral consequence of rapid sensory adaptation that permits iL3s to flexibly adjust their host-seeking behaviors after encountering a non-host heat source.

## The *Ss*-AFD-rGCs each confer sensitivity to temperature ranges spanning ambient and host body heat

Next, we sought to identify molecular mechanisms that underlie the expanded responsiveness of *Ss*-AFD to mammalian body temperatures. In *C. elegans*, two of the AFD-specific rGCs, *Ce-*GCY-18 and *Ce-*GCY-23, act as thermoreceptors^35^. Specifically, misexpression of *Ce*-GCY-18 and *Ce*-GCY-23, but not *Ce*-GCY-8, can confer TAX-4-dependent sensitivity to thermal stimuli^35^. We wanted to test whether the *S. stercoralis* AFD-rGCs (*Ss*-GCY-23.1, *Ss*-GCY-23.2, and *Ss*-GCY-23.3) are thermoreceptors, and if so whether they display response properties that might contribute to the expanded range of *Ss*-AFD responsiveness. We therefore ectopically expressed each *Ss*-AFD-rGC in *C. elegans* ASE chemosensory neurons also expressing the YC3.60 calcium sensor, then measured responses to naturalistic warming temperature ramps*. Ce*-ASE was chosen as the ectopic expression site, rather than *Ce*-AFD, in order to examine the thermosensory properties of the *Ss*-AFD-rGCs independent of mechanisms for *Ce*-AFD-specific signal amplification^26,35^.

We found that each *Ss*-AFD-rGC is capable of conferring thermosensitivity in *Ce*-ASE, and that their functional properties differ from each other and from those of the *C. elegans* AFD-rGCs (Fig. 4a,b, Fig. S10). For *Ss*-GCY-23.1 and *Ss*-GCY-23.3 ectopic expression, the threshold (T*_ASE_) of temperature-evoked activity is near mammalian skin temperature; *Ss*-GCY-23.2 expression results in thermosensitive activity with a T*_ASE_ near ambient cultivation temperature (Fig. 4c). Notably, all three *Ss*-AFD-rGCs confer thermosensitivity up to host body temperatures. Across the *Ss*-AFD-rGCs, the temperature driving maximal responses varies such that *Ss*-GCY-23.3 responses peak at warmer temperatures than *Ss*-GCY-23.1 and *Ss*-GCY-23.2 (Fig. 4c). These results suggest that changes in the functional properties of *Ss*-AFD-rGCs, including an expanded responsiveness to mammalian body temperatures relative to *C. elegans* homologs, may contribute to both parasite-specific encoding of thermal cues in *Ss*-AFD and the preference of *S. stercoralis* iL3s for mammalian body heat.

**Figure 4.**
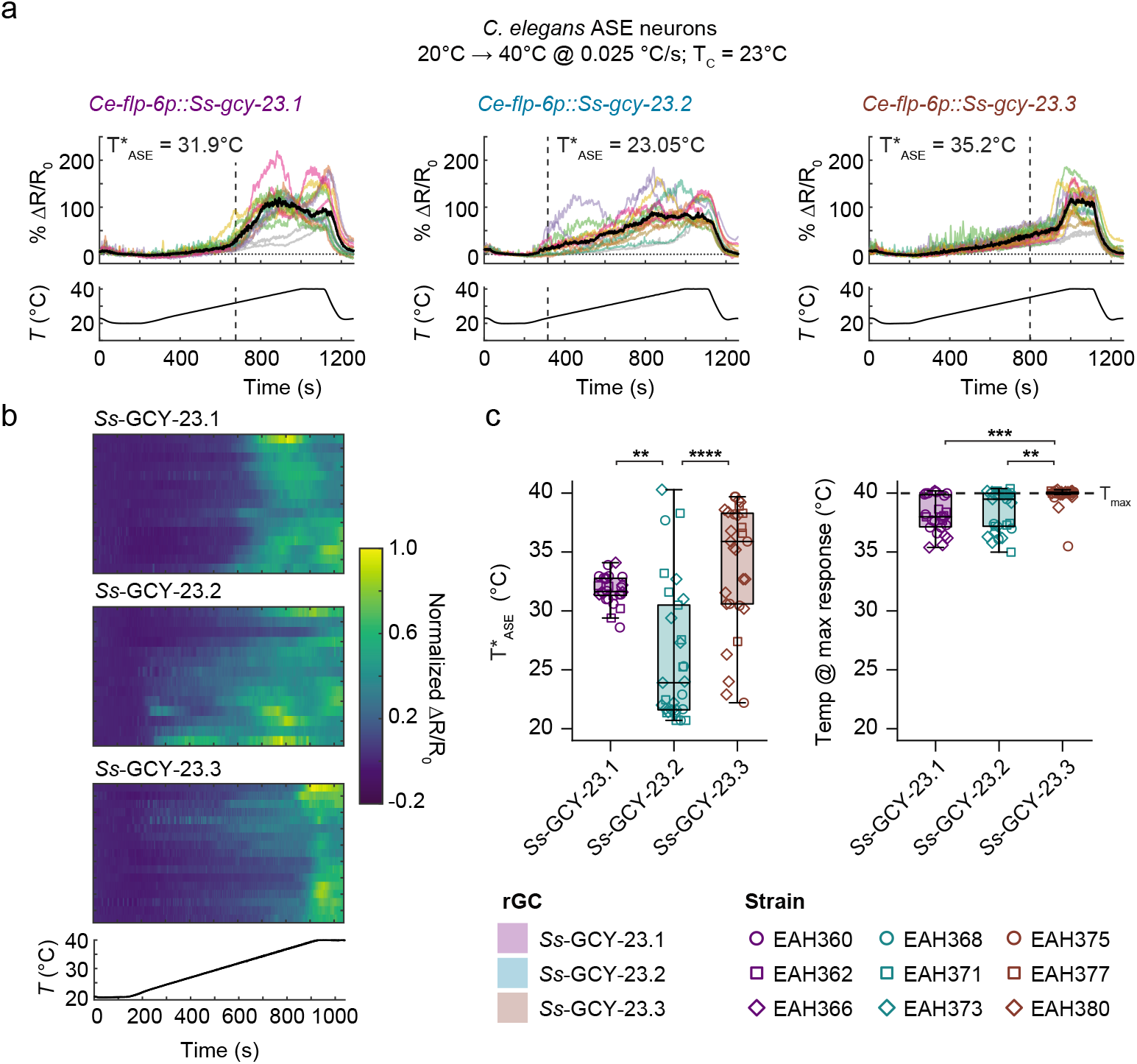
All three *S. stercoralis* AFD-specific rGCs are thermoreceptors. **a.** Responses of *C. elegans* ASER/L neurons expressing YC3.60 and *Ss*-GCY-23.1 (left), *Ss*-GCY-23.2 (middle), or *Ss*-GCY-23.3 (right) to naturalistic warming temperature ramps. Warming stimulus ramp: 20-40°C at 0.025 °C/s. Vertical dashed lines indicate average thermal threshold (T*_ASE_), defined as the first temperature where ΔR/R_0_ deviates from the average *Ce*-ASE_WT_ response by at least 3 * STD of ASE_WT_ R/R_0_ for a minimum of 40 seconds. Colored traces indicate responses of individual worms that crossed the thermal threshold; gray traces indicate responses that did not. Black traces show median responses; only worms with a response above threshold are included in averages. Temperature traces show the average ± SD recorded temperature (lines + shading; in some cases shading is too small to be visible). R_0_ (horizontal dotted lines) = response at 20°C. T_C_ = 23°C. n = 15 worms above threshold out of 17 total worms (*Ss*-GCY-23.1), 14 of 15 worms (*Ss*-GCY-23.2), and 17 of 19 worms (*Ss*-GCY-23.3). **b.** Heatmaps of individual *C. elegans* ASER/L responses from panel **a**. Responses are normalized to the maximum ΔR/R_0_ response for each strain, with R_0_ set to zero. Responses are ordered by hierarchical cluster analysis. **c.** Thermal thresholds (left) and temperatures eliciting highest calcium responses (right) for *C. elegans* ASE neurons expressing *Ss-gcy-23.1* (purple icons), *Ss-gcy-23.2* (teal icons), and *Ss-gcy-23.3* (red icons) measured in individual worms from three replicate strains (n = 5-17 worms per strain). Only worms with a response crossing the thermal threshold are included in quantifications. Boxes show medians and interquartile ranges, whiskers show min and max values. Horizontal dashed line indicates T_max_ (40°C). n = 5-17 worms per strain, with 26, 29, and 29 worms per genotype (*Ss*-GCY-23.1, *Ss*-GCY-23.2, and *Ss*-GCY-23.3, respectively). \*\**p*<0.01, \*\*\**p*<0.001, \*\*\*\**p*<0.0001, two-sided Kruskal-Wallis test with Dunn’s post-test; multiplicity adjusted *p* values are reported.

## Discussion

Here, we identify multiple parasite-specific molecular, cellular, and behavioral adaptations in the thermosensory circuit of the human parasite *S. stercoralis*. We present the first uses of single-cell genetic targeting and neural imaging methods in a non-*Caenorhabditis* nematode, as well as the first direct demonstration of parasite specific neural encoding strategies that underlie sensory-driven host seeking by soil-transmitted parasitic nematodes. Our results provide mechanistic insight into the neural basis of heat seeking, an essential component of the ability of soil-transmitted parasitic nematodes to target human hosts. Historically, most research on parasitic nematodes has used the free-living nematode *Caenorhabditis elegans* as a genetically-tractable model for human parasites. Our data indicate that this approach is not sufficient, and emphasize the essential role of mechanistic studies in parasitic nematodes themselves for understanding the neural basis of parasitism in these species. Parasitic nematode infections are one of the world’s most neglected sources of chronic disease and economic burden; parasitic sensory circuits are a promising, yet largely unexplored, target for intervention.

## Methods

All protocols and procedures involving vertebrate animals were approved by the UCLA Office of Animal Research Oversight (Protocol 2011-060-AM-003), which adheres to the standards of the AAALAC and the *Guide for the Care and Use of Laboratory Animals*.

### Molecular Biology

Identification of the following *S. stercoralis* and *S. ratti* genes was based on protein sequence homology with *C. elegans* GCY-8, GCY-18, and GCY-23: *Ss-gcy-23.1* (SSTP_0000354300), *Ss-gcy-23.2* (SSTP_0000775800)*, Ss-gcy-23.3* (SSTP_0000846800)*, Sr-gcy-23.1* (SRAE_2000430600)*, Sr-gcy-23.2* (SRAE_0000007100), and *Sr-gcy-23.3* (SRAE_X000020900). Amino acid sequences were retrieved from WormBase ParaSite^52^. Pairwise alignment of amino acid sequences was calculated in Geneious Prime (v2020.1.1) using global alignment with free end gaps and a Blosum62 cost matrix. Tree construction was performed in Geneious Prime using a Jukes-Cantor model with neighbor-joining.

Promoter sequences for all *S. stercoralis* genes and *Sr-gcy-23.2* were extracted by PCR-amplifying ∼1.7-2 kb upstream of the predicted first exons; some sequences include portions of the first exon. Primers used are listed in Supplemental Data File 2. Putative promoter sequences were cloned into either the *Strongyloides* expression vector pPV254 (containing *Ss-act-2p::GFP*; a gift from Dr. James Lok, UPenn) in place of the *Ss-act-2* promoter, or into a *Strongyloides* promoter-less expression vector (pASB10, which features a multiple cloning site and the *Ss-era-1* 3’UTR^39^) along with *wrmScarlet-I*^53^ (obtained from Dr. Hillel Schwartz, Caltech). The following constructs were generated: *Ss-gcy-23.1p::GFP* (pASB51)*, Ss-gcy-23.2p::GFP* (pASB41)*, Ss-gcy-23.3p::GFP* (pASB42)*, Sr-gcy-23.2p::GFP* (pJHL09)*, and Sr-gcy-23.2p::wrmScarlet-I* (pASB25).

For expression of HisCl1 in *S. stercoralis* AFD, we first calculated a *Strongyloides* codon-optimized *HisCl1* sequence (*strHisCl1*) by hand^42,54^. To generate a *Sr-gcy-23.2p::strHisCl1* construct (pASB30), *strHisCl1* was synthesized by Genewiz (South Plainfield, NJ) and cloned into pASB10 with the *Sr*-*gcy-23.2* promoter from pJHL09. For expression of the calcium sensor YC3.60^43^ in *S. stercoralis* AFD, we calculated a *Strongyloides* codon-optimized *YC3.60* sequence (*strYC3.60*) by hand. To generate a *Sr-gcy-23.2p::strYC3.60* construct (pASB52), *strYC3.60* was synthetized by Genewiz and cloned into pASB30, replacing *strHisCl1*.

For ectopic expression of *S. stercoralis* AFD-rGCs in *C. elegans* ASE neurons, cDNA sequences were identified based on gene predictions in WormBase ParaSite^52,55^. Sequences were codon-optimized for *C. elegans* and 2 synthetic introns were inserted into the sequence using the *C. elegans* Codon Adapter^56^. To generate a *flp-6p::Ss-gcy-23.1::SL2::mCherry* construct (pASB57), the cDNA sequence was synthesized by Genewiz and used to modify the plasmid AT1_66, obtained from Dr. Piali Sengupta (Brandeis)^35^. To generate *flp-6p::Ss-gcy-23.2::SL2::mCherry* (pASB61) and *flp-6p::Ss-gcy-23.3::SL2::mCherry* (pASB62) constructs, cDNA sequences were synthesized and inserted into AT1_66 by GenScript (Piscataway, NJ). A full list of all plasmids used in this study are provided in Supplemental Data File 2.

### Strains

*S. stercoralis* (UPD strain) was maintained as previously described^10^. Transgenic *S. stercoralis* F_1_ iL3s were generated by microinjection of plasmid constructs into free-living adult females using standard methods^39^. Fecal-charcoal plates containing microinjected females and wild-type adult males were incubated at 23°C for a minimum of 5 days prior to screening and experimental use. To generate transgenic F_1_ iL3s expressing fluorescent reporters, plasmids (including: pJHL09, pASB25, pASB41, pASB42, pASB51, and pMLC41^10^) were microinjected at 40-100 ng/μL (Supplemental Data File 2). Targeted mutagenesis of *Ss-tax-4* was accomplished using established *Strongyloides* CRISPR/Cas9 protocols^10,47^, with the following modifications. For generating *Ss-tax-4* iL3s, injection mixes contained 80 ng/μL pMLC47, 80 ng/μL pEY11, and 40 ng/μL pPV540. For generating no-Cas9 controls, injection mixes contained 80 ng/μL pMLC47 and 80 ng/μL pEY11. For calcium imaging of *S. stercoralis* AFD, *Sr-gcy-23.2p::strYC3.60* F_1_ iL3s were generated by microinjecting pASB52 at a concentration of 80 ng/μL. For generating *Sr-gcy-23.2p::strHisCl1* iL3s, injection mixes contained 80 or 100 ng/μL pASB30 with 50 ng/μL pJHL09 as a co-injection marker.

*C. elegans* adults were raised on 2% Nematode Growth Media (NGM) plates seeded with a lawn of *Escherichia coli* OP50 bacteria, as per standard methods^57^. *C. elegans* experiments exclusively used young adult hermaphrodites; adult males were used only for crosses. Calcium imaging of *C. elegans* AFD was performed using strain IK890 (*njEx358[gcy-8::YC3.60, ges-1::NLS-tagRFP]*), which was obtained from Dr. Ikue Mori (Nagoya University). Calcium imaging of *C. elegans* ASE was performed using strain XL115 *lin-15(n765) ntIs14[flp-6::YC3.60; lin-15+]*, which was obtained from Dr. Shawn Lockery (U. Oregon). Independent strains for ectopic expression of *S. stercoralis* AFD-rGCs in *C. elegans* ASE (Supplemental Data File 2, strains EAH359-EAH379) were generated by microinjecting test plasmids (pASB57, pASB61, pASB62) into N2 hermaphrodites at 20 ng/μL. Each ectopic expression strain was crossed to XL115 to generate the strains used for calcium imaging (Supplemental Data File 2, strains EAH360-EAH380).

### Fluorescent Microscopy

For fluorescent microscopy, the F_1_ iL3 progeny of microinjected females were recovered and screened for GFP or wrmScarlet-I expression on a Leica M165 FC fluorescence microscope using a previously established nicotine-paralysis method^47^. Transgenic iL3s were transferred to a small watch glass containing BU saline^58^ and allowed to recover overnight. Animals were exposed to 50-100 mM levamisole in BU saline, then mounted on a slide with 5% Noble agar dissolved in BU saline, coverslipped, and sealed with quick-drying nail polish. Epifluorescence and DIC images were taken using a Zeiss AxioImager A2 microscope with an attached Hamamatsu ORCA-Flash camera (C13440-20CU-KIT Flash4.0 V3 sCMOS). Confocal images were acquired using a Zeiss LSM880 confocal microscope (Broad Stem Cell Research Center, Molecular, Cellular, and Developmental Biology Microscopy Core Facility, UCLA). Images were processed using the Zeiss Zen pro software (v2.6). Image montages were generated using FIJI.

### Behavioral Assays

Thermotaxis assays were performed using a custom thermoelectric behavioral arena^10^. For experiments quantifying the distribution of a population, worms were monitored using one or two 5 mega-pixel CMOS cameras (BTE-BO50-U, Mightex Systems) at a frame rate of at least 1 frame per minute. The population-level movement of worms was analyzed as previously described^10^. For worm tracking experiments, worm movements were imaged at a frame rate of 0.5 frames per second, and individual iL3 trajectories were quantified *post hoc* as previously described^10^, using updated custom MATLAB scripts.

For testing wild-type thermotaxis behaviors across different cultivation temperatures, iL3s were collected from 7-14 day old fecal-charcoal plates using a Baermann apparatus^59^ or removed directly from the fecal charcoal lid and then stored in a watch glass containing BU saline at either 15°C or 23°C for at least two hours. For negative thermotaxis behaviors, iL3s were placed in a 12-22°C gradient (T_start_ = 14°C, 17°C, or 20°C) for 30 minutes. For positive thermotaxis behaviors, iL3s were placed in a ∼21-33°C gradient (T_start_ = 23°C) for 20 minutes. The behavioral data for unstimulated wild-type iL3s (Fig. S2b) are from Bryant *et al* 2018^10^.

For measuring reversal behaviors, 7-14 day old fecal-charcoal plates were incubated for at least two hours at 15°C or 23°C. iL3s were collected using a Baermann apparatus or removed directly from the lid of the fecal-charcoal plate and stored in a watch glass containing BU saline at their cultivation temperature for at least two hours. Two thermal gradients were used to elicit reversal behaviors: a 12-22°C gradient (T_start_ = 20°C) and a 15-25°C gradient (T_start_ = 23°C). For testing the percentage of the population that engaged in reversal behaviors, at least 150 iL3s were placed on a thermotaxis plate and their migration was imaged for 20-30 minutes. For tracking individual iL3 trajectories during reversal behaviors, either single iL3s or a small group (<100) of iL3s were placed on a thermotaxis plate and their migration was imaged for 20 min.

For testing negative thermotaxis by *Ss-tax-4* iL3s, either *Ss-tax-4* iL3s or no-Cas9 control iL3s were screened for mRFPmars expression as previously described^10,47^. 1-2 mRFPmars-expressing *S. stercoralis* iL3s were placed at 20°C in a ∼17-26°C gradient for 15 min. Following the cessation of recording, single iL3s were recollected for individual genomic DNA preparations. mRFPmars-positive iL3s in which *Ss-tax-4* expression was fully disrupted either by deletion or integration events (*i.e*., *Ss-tax-4* iL3s) were identified by *post-hoc* single-worm genotyping, as previously described^10,47^. Trajectories of *Ss-tax-4* iL3s were tracked and quantified following *post-hoc* genotyping.

For testing the effect of HisCl1-mediated AFD silencing on positive and negative thermotaxis, *Sr-gcy-23.2p::strHisCl1 + Sr-gcy-23.2p::GFP* iL3s were screened for GFP expression as follows: F_1_ iL3 progeny of microinjected females were recovered from fecal-charcoal cultures using a Baermann apparatus and stored in a small watch glass containing 2-3 mL BU saline. ∼10 μL of iL3-containing BU saline was placed on a 2% NGM plate containing a lawn of OP50, and migrating iL3s were screened at low magnification for strong GFP expression. Recovered iL3s were evenly split into two watch glasses containing either BU saline or BU saline + either 20 or 50 mM histamine. Watch glasses were incubated at 23°C for at least 2 h. For behavioral experiments, iL3s were transferred in batches (<20 worms per experiment) to 3% thermotaxis agar plates^10,60^ with or without 10 mM histamine. For positive thermotaxis assays, iL3s were placed at 30°C in an ∼22-34°C gradient; for negative thermotaxis assays, iL3s were placed at 20°C in a ∼17-26°C gradient. For both positive and negative thermotaxis, assay duration was 10 min. On individual experimental days, behavioral assays were always conducted with paired histamine-positive and no-histamine control experiments. Furthermore, the researcher performing the assay was fully blinded to the experimental condition: blinding was implemented as recovered GFP-positive iL3s were split into BU saline or BU saline + histamine watch glasses (*i.e*., the researcher splitting the population was unaware of which watch glass contained histamine-spiked BU saline). Blinding was lifted only following *post-hoc* tracking.

### Calcium Imaging

For calcium imaging of *Ss*-AFD, the F_1_ iL3 progeny of microinjected females were recovered and screened for the presence of YC3.60 using methods previously established for fluorescent screening of nicotine-paralyzed transgenic iL3s^47^. Prior to calcium imaging, *C. elegans* adults on 2% NGM plates with an OP50 lawn or *S. stercoralis* iL3s in BU saline-containing watch glasses were incubated overnight at either 23°C or 15°C. Animals were exposed to 100 mM levamisole in BU saline (*S. stercoralis* iL3s) or 10 mM levamisole in M9 saline (*C. elegans* adults), then mounted on a slide with 5% Noble agar dissolved in BU or M9 saline, coverslipped, and sealed with quick-drying nail polish.

Calcium imaging was performed using a Zeiss AxioImager A2 microscope equipped with a 78 HE ms CFP/YFP dual camera filter set, a Hamamatsu W-View Gemini image splitter, and a Hamamatsu ORCA-Flash camera for simultaneous acquisition of CFP and YFP images. Images were acquired every 250 ms using the Zeiss ZEN software with a time lapse module.

Temperature ramps were delivered to slide-mounted worms using a custom thermal stimulator (Fig. S3a) based on established *C. elegans* systems^46^. An annular-style Peltier element (430533-502, Laird Thermal Systems) was mounted on a custom aluminum heat block held on the microscope stage with a custom 3D-printed stage clamp. The custom heat block was attached to a temperature-controlled recirculating water bath (13-874-180, Fisher Scientific) set to the cultivation temperature. The Peltier element was controlled by a closed-loop control circuit consisting of a PID controller (ATEC302, Accuthermo Technology), an H-bridge amplifier (FTX700D, Accuthermo Technology), two 12 V, 30 W AC/DC power converters (LS35-12, TDK-Lambda Americas Inc), and a 10 kΩ thermistor (USP12837, Littelfuse) that was attached to the coverslip near the immobilized worm with adhesive tape. Slides containing worms were placed on the Peltier element; halocarbon oil was placed between the Peltier element and the glass slide to ensure strong thermal transfer. Slides were anchored to the aluminum heat block using adhesive tape. Prior to the start of imaging, slides were brought to cultivation temperature (23°C or 15°C) for 5 min using the ATEC302 software general control mode. During imaging, temperature ramps were delivered using the ATEC302 software programmable step temperature control function. A full list of all thermal ramps used in this study is provided in Supplemental Data File 2. Measured temperatures at the coverslip surface were recorded using the ATEC302 software data log function.

From time-lapse dual-wavelength images containing aligned CFP and YFP signals we calculated the mean intensity over time for soma and background regions of interest (ROI) using the ZEN software. CSV files containing imaging data and temperature log files were saved, then processed using custom MATLAB scripts. Briefly, CFP and YFP signals from soma and background ROIs were used to calculate the baseline-corrected YFP/CFP ratio (% ΔR/R_0_); YFP signals were corrected for bleed-through from the CFP channel. Calcium responses were aligned relative to the ATEC302 temperature log based on file time stamps. Plots of aligned calcium responses and temperature records were saved as EPS files. For heatmaps, individual calcium traces (heatmap rows) were ordered using average linkage clustering; heatmap colors were defined using the viridis color map.

Custom MATLAB code was used to calculate: linear/monotonic regressions at different temperature windows, thermal threshold (either T*_AFD_ or T*_ASE_), temperatures eliciting maximal and minimal calcium responses, and the response magnitude at the hottest stimulus temperature (T_max_) or the coolest stimulus temperature (T_min_), across different time windows. For AFD recordings, thermal threshold (T*_AFD_) is defined as the first temperature where the absolute value of the AFD response is at least 3*STD from the response at T_0_ for the amount of time it takes the warming/cooling temperature ramp to change 0.25°C (either 2.5 or 10 s). For *C. elegans* ectopic expression recordings, thermal threshold (T*_ASE_) was defined as the first temperature where the absolute value of the *Ce*-ASE_experimental_ response deviates from the average *Ce*-ASE_WT_ response by at least 3*STD of the *Ce*-ASE_WT_ response for a minimum of 40 seconds.

### Quantification and Statistical Analysis

Statistical analyses were conducted using GraphPad Prism 9. Exact *p* values for all statistical tests are provided in Supplemental Data File 1. Power analyses to determine appropriate sample sizes were conducted using G*Power 3.1. For plotting single-worm tracking quantifications and calcium imaging quantifications, data were imported into R v3.6.3 and plotted using the ggplot2 package v3.3.2^61^.

## Data Availability

The data that support the findings of this study are available on GitHub (https://github.com/HallemLab/Bryant_et_al_2021).

## Code Availability

Source code for the following tasks is available on GitHub: acquisition, analysis, and plotting of single worm tracks; preprocessing calcium imaging data; and plotting quantifications from worm tracking and calcium imaging experiments (https://github.com/HallemLab/Bryant_et_al_2021). Parts lists and wiring diagrams for custom hardware are also available in the same GitHub repository.

## Acknowledgements

We gratefully acknowledge Dirk Williams and the UCLA R&D Shops, as well as the UCLA MCDB/BSCRC microscopy core. We thank Dr. Josh Hawk for advice related to the design of thermal stimulators for calcium imaging. We also thank Dr. Piali Sengupta for advice related to the naming of *Strongyloides* AFD-rGCs. We thank Dr. Pavak Shah, Dr. Michelle Castelletto, and Dr. Ruhi Patel for insightful comments on the manuscript. This work was funded by an A.P. Giannini Postdoctoral Fellowship (A.S.B.); and a Burroughs-Wellcome Fund Investigators in the Pathogenesis of Disease Award, HHMI Faculty Scholar Award, and NIH R01AI136976 (E.A.H.).

## Author Contributions

A.S.B. and E.A.H. conceived the study and wrote the manuscript. A.S.B., F.R., and J.L. performed all experiments and analyzed the data. All authors read and approved the final manuscript.

