## Supplemental Material for "The neural basis of temperature-driven host seeking in the human threadworm *Strongyloides stercoralis*"

**Supplemental Figure Legends**

**Figure S1. Genetic identification of the *S. stercoralis* AFD neuron pair. a.** Amino acid sequence homology of the *C. elegans* AFD-rGCs (*Ce*-GCY-8, *Ce*-GCY-18, and *Ce*-GCY-23) and the *S. stercoralis* AFD-rGCs (left) or the *S. ratti* AFD-rGCs (right). Individual tables show % sequence identity between the full-length proteins. Bold values indicate the highest % sequence identity for each parasite protein. **b.** The promoter region of *Sr-gcy-23.2* drives wrmScarlet-I expression in *S. stercoralis* iL3s that co-localizes with expression of the three endogenous *S. stercoralis* AFD rGCs (left: *Ss-gcy-23.1*; middle: *Ss-gcy-23.2*; right: *Ss-gcy-23.3*). Scale bar = 20 µm. Anterior is left. **c.** Expression of *Sr*-*gcy-23.2* co-localizes with expression of *Ss-tax-4.* Star indicates the soma of a co-labeled *Ss*-AFD neuron. Arrowhead indicates another *Ss-tax-4*+ neuron that does not express the *Ss*-AFD-specific marker. Scale bar = 10 µm. Anterior is left, dorsal is up.

**Figure S2. Chemogenetic silencing of *S. stercoralis* AFD. a.** Workflow for testing the effect of inducible HisCl1-mediated silencing of *S. stercoralis* AFD on the behavior of *S. stercoralis* iL3s. DNA plasmids containing a *Strongyloides* codon-optimized HisCl1 gene (*strHisCl1*) and *GFP* under the control of the *Sr-gcy-23.2* promoter are co-injected into the gonads of free-living adult females. The F_1_ progeny are screened for the presence of GFP. GFP*+* F_1_ iL3s are divided into two groups, and incubated for at least 2 h in BU saline or BU saline + 20 or 50 mM histamine. Worms are then batch-tested in thermotaxis assays (<20 worms per experiment). All steps following initial GFP screening, including analysis of worm tracks, are performed while the experimenter is blinded to the assay conditions (BU vs. histamine). **b.** Chemogenetic silencing of *S. stercoralis* AFD does not affect worm motility. Comparison of the mean speed of unstimulated wild-type iL3s (gray squares) placed on an isothermal room temperature plate versus *Sr-gcy-23.2p::strHisCl1*; *Sr-gcy-23.2p::GFP* iL3s treated with BU saline or histamine (gray and yellow circles, respectively) placed in a ~22-34°C gradient for 10 min. T_start_ = 30°C. n = 15 worms for wild-type iL3s, 38 worms for *Sr-gcy-23.2p::strHisCl1*; *Sr-gcy-23.2p::GFP* iL3s treated with BU saline, and 36 worms for *Sr-gcy-23.2p::strHisCl1*; *Sr-gcy-23.2p::GFP* iL3s treated with histamine. ns = not significant (*p*>0.05), *****p*<0.0001, two-sided Kruskal-Wallis test with Dunn’s multiple comparisons tests; multiplicity adjusted *p* values are reported. *Sr-gcy-23.2p::strHisCl1*; *Sr-gcy-23.2p::GFP* iL3 data are replotted from Figure 1. Unstimulated wild-type iL3 data are from Bryant *et al* 2018^10^.

**Figure S3. Thermal stimulation of immobilized *C. elegans* adults and *S. stercoralis* iL3s. a.** Schematic of the custom-built thermoelectrical control system capable of delivering precisely controlled thermal stimuli to immobilized worms during calcium imaging with a Zeiss AxioImager A2 microscope. The use of an annular-style Peltier element permits locating worms in the field of view using transmitted light. **b.** Diagram of the naturalistic thermal stimulus that mimics the average trajectories of *S. stercoralis* iL3s engaging in positive thermotaxis in a ~22-34°C thermal gradient (cultivation temperature, T_C_ = 23°C). Worms are held at their cultivation temperature under the microscope for 5 min prior to the start of imaging (dashed line). Warming ramp slope: 0.025 °C/s. **c.** Schematic of the naturalistic thermal stimulus used for imaging the activity of worms cultivated at 15°C. Timing and slope of temperature stimulus is as shown in panel **b**, with absolute temperatures shifted down by 8°C.

**Figure S4. Response of *Ss*-AFD to a naturalistic warming stimulus spanning ambient and core body temperatures. a.** *S. stercoralis* AFD responses to a 20-40°C warming temperature ramp. Colored traces show the responses of individual worms; black traces show median responses. **b.** Heatmaps of individual *Ss*-AFD responses shown in panel **a**. Heatmaps show responses normalized to the maximum ΔR/R_0_ response, with R_0_ set to zero. Responses are ordered by hierarchical cluster analysis. For all plots, temperature traces show the average recorded temperature (mean in solid line, SD in shading; in some cases, shading is too small to be visible). Warming ramp slope = 0.025 °C/s. R_0_ (horizontal dotted lines) = response at 20°C. n = 15 worms. T_C_ = 23°C.

**Figure S5. *Ss*-AFD response to rapid warming temperature ramps. a.** Diagram of a rapid warming stimulus spanning a range of temperatures that elicits strong thermotaxis migration in *S. stercoralis* iL3s (17-40°C). T_C_ = 23°C. Warming ramp slope: 0.1 °C/s. Worms are held at T_C_ under the microscope for 5 min prior to the start of imaging (dashed line). **b*.*** *C. elegans* AFD responses to a rapid 17-40°C warming temperature ramp. Colored traces show the responses of individual worms; black traces show median responses. Temperature traces show the average recorded temperature (mean in solid line, SD in shading; shading is too small to be visible). Horizontal dotted line indicates baseline (R_0_) response at 17°C. Heatmap shows responses normalized to the maximum *Ce*-AFD ΔR/R_0_ response, with R_0_ set to zero. Responses are ordered by hierarchical cluster analysis. **c.** *S. stercoralis* AFD responses to a rapid 17-40°C warming temperature ramp. *Left:* Traces in which the temperature eliciting the minimum *Ss*-AFD response is greater than the cultivation temperature (T_C_). *Middle*: Traces in which the temperature eliciting the minimum *Ss*-AFD response is less than or equal to T_C_. *Right*: Heatmaps show responses normalized to the second largest *Ss*-AFD ΔR/R_0_ response, with R_0_ set to zero; the largest response is saturated. All other conventions are as in panel **b**. **d.** Temperature that elicits the lowest calcium response in *Ce*-AFD and *Ss*-AFD, in response to a rapid 17-40°C warming ramp. Responses are categorized based on whether the temperature eliciting the minimum response is less than/equal to T_C_ (n = 12 neurons for *S. stercoralis*, 20 for *C. elegans*) or greater than T_C_ (n = 7 *S. stercoralis* neurons). Horizontal dashed lines indicate T_C_ (23°C) and T_min_ (17°C). **p*<0.05, ***p*<0.01, *****p*<0.0001, Kruskal-Wallis test with Dunn’s multiple comparisons test. **e.** Proportion of *Ce*-AFD and *Ss*-AFD recordings in which the temperature eliciting the minimum response is greater (teal) or less than/equal to T_C_ (gray), for both naturalistic and rapid ramp speeds. The proportion of *Ss*-AFD recordings that exhibit a significant warming-triggered inhibition was greater in response to slower temperature ramps, versus fast temperature ramps (*p*<0.005, Fisher’s exact test). n = 19 neurons from 18 worms (*Ss*-AFD rapid ramp), 20 neurons from 20 worms (*Ce*-AFD rapid and naturalistic ramps), 30 neurons from 30 worms (*Ss*-AFD naturalistic ramp)*.* **f.** Thermal threshold (T*) of *C. elegans* AFD and *S. stercoralis* AFD calcium responses at two ramp speeds. T*_AFD_ is defined as the first temperature where % ΔR/R_0_ is significantly different from baseline (3 * STD R_0_) for the time required for the temperature to increase 0.25°C (either 2.5 or 10 seconds). Data are responses to either naturalistic warming ramps (20-34°C (circles) or 20-40°C (squares) at 0.025 °C/s) or fast warming ramps (17-40°C (diamonds) at 0.1 °C/s). ns = not significant (*p*>0.05), two-way ANOVA with Šidák’s multiple comparisons test. n = 19 neurons from 18 worms (*Ss*-AFD rapid ramp), 20 neurons from 20 worms (*Ce*-AFD rapid and naturalistic ramps), 30 neurons from 30 worms (*Ss*-AFD naturalistic ramp). **g.** Quantification of linear correlation between temperature and calcium responses at 25°C and higher. Values are Pearson’s correlation coefficients, icon conventions are the same as panel **f**. ns = not significant (*p*>0.05), **p*<0.05, *****p*<0.0001, two-way ANOVA with Šidák’s multiple comparisons test. **h.** Temperature that elicits the highest calcium response in *Ce*-AFD and *Ss*-AFD, in response to a rapid 17-40°C warming ramp. Horizontal dashed line indicates T_max_ (40°C). ***p*<0.005, Mann-Whitney test. For **f-g,** naturalistic ramp data are replotted from Figure 2. For **d** and **f-h**, boxes show medians and interquartile ranges, whiskers show min and max. Unless otherwise specified, n = 19 neurons from 18 worms (*S. stercoralis*), 20 neurons from 20 worms (*C. elegans*). All statistical tests are two-sided; multiplicity adjusted *p* values are reported when appropriate.

**Figure S6. Exposure to a new cultivation temperature shifts the *Ss*-AFD thermal threshold. a.** *Ce*-AFD responses to a 12-26°C temperature ramp, after overnight cultivation at 15°C. *Left:* Colored traces show the responses of individual worms; black traces show the median response. Horizontal dotted line indicates the baseline (R_0_) response at 12°C. *Right:* Heatmap shows responses normalized to the maximum *Ce*-AFD ΔR/R_0_ response, with R_0_ set to zero. Temperature traces show the average recorded temperature (mean in solid line, SD in shading; in some cases, shading is too small to be visible). Warming ramp slope = 0.025 °C/s. R_0_ (horizontal dotted lines) = response at 12°C. n = 16 worms. **b.** *Ss*-AFD responses to a 12-26°C temperature ramp, after overnight cultivation at 15°C. Heatmap shows responses normalized to the second largest *Ss-*AFD ΔR/R_0_ response, with R_0_ set to zero; the largest response is saturated. All other conventions are as in panel **a**. n = 17 worms. **c.** *Ss*-AFD responses to a 12-32°C temperature ramp, after overnight cultivation at 15°C. Heatmap shows responses normalized to the maximum *Ss-*AFD ΔR/R_0_ response, with R_0_ set to zero. All other conventions are as in panel **a**. n = 16 worms. **d.** Effect of cultivation temperature (15°C versus 23°C) on *Ce-*AFD and *Ss-*AFD thermal threshold (T*_AFD_). Data are responses to ramps with a net temperature change of 14°C (circles) or 20°C (squares). n = 16 worms (*Ce*-AFD, T_C_ = 15°C), 20 worms (*Ce*-AFD, T_C_ = 23°C), 33 worms (*Ss*-AFD, T_C_ = 15°C), 30 worms (*Ss*-AFD, T_C_ = 23°C). *****p*<0.0001, 2-way ANOVA with Tukey’s multiple comparisons tests. Responses for T_C_ = 23°C are re-plotted from Figure 2. **e.** Temperature that elicits the lowest calcium response in *Ce*-AFD and *Ss*-AFD, in response to either a 12-26°C (circles) or a 12-32°C (squares) warming ramp. T_C_ = 15°C. Horizontal dashed lines indicate T_C_ (15°C) and T_min_ (12°C). n = 16 worms (*Ce*-AFD), 33 worms (*Ss*-AFD). *****p*<0.0001, two-sided Mann-Whitney test. **f.** Temperature that elicits the highest calcium response in *Ce*-AFD and *Ss*-AFD, in response to a 12-26°C warming ramp. T_C_ = 15°C. Horizontal dashed line indicates T_max_ (26°C). n = 16 worms (*Ce*-AFD), 17 worms (*Ss*-AFD). ****p*<0.001, two-sided Mann-Whitney test. For all plots: icons indicate individual worms, boxes show median and interquartile range, whiskers show min and max values.

**Figure S7. *S. stercoralis* iL3s do not display a single preferred cold temperature. a.** *Left:* Representative distribution of *S. stercoralis* iL3s cultivated at 23°C and then placed at 14°C, 17°C, or 20°C in a 12-22°C gradient for 30 min. Colored inverted triangles indicate the different possible T_start_ locations. iL3s display negative thermotaxis in all conditions. The final temperature reached varies with T_start_, suggesting that *S. stercoralis* iL3s do not have a single preferred low temperature. *Right*: Median final distribution of *S. stercoralis* iL3s placed at 14°C (blue triangles), 17°C (yellow circles) or 20°C (red squares) in a 12-22°C gradient. Colored inverted triangles indicate T_start_. Graphs show medians and interquartile ranges. T_C_ = 23°C; duration: 30 min; n = 15 trials for each condition with >50 iL3s per trial. **b.** *Left:* Representative distribution of *S. stercoralis* iL3s cultivated at 15°C for 2-8 h and then placed at 14°C, 17°C, or 20°C in a 12-22°C gradient for 30 min. iL3s display negative thermotaxis, with a T_C_-dependent shift in the range of preferred low temperatures. *Right*: Median final distribution of *S. stercoralis* iL3s placed at 14°C (blue triangles), 17°C (yellow circles) or 20°C (red squares) in a 12-22°C gradient. Colored inverted triangles indicate T_start_. Graphs show medians and interquartile ranges. T_C_ = 15°C; duration: 30 min; n = 16 trials (14°C), 17 trials (17°C), 15 trials (20°C). In all cases >50 iL3s were used per trial. **c.** Median final distribution of *S. stercoralis* iL3s placed at 14°C (left), 17°C (middle), or 20°C (right) in a 12-22°C gradient. Green lines and icons indicate worms cultivated at 15°C for 2-8 h. Purple lines and icons indicate worms cultivated at 23°C. Colored inverted triangles indicate T_start_, as in panels **a**-**b**. Graphs show medians and interquartile ranges. Data are replotted from panels **a**-**b;** n values are as indicated in panels **a**-**b**. **d.** Comparison of the average final temperature reached by *S. stercoralis* iL3s, across starting temperatures and cultivation temperatures. Blue icons are worms placed at 14°C, yellow icons are worms placed at 17°C, and red icons are worms placed at 20°C. Circles are worms cultivated at 23°C, squares are worms cultivated at 15°C. ***p*<0.01, *****p*<0.0001, 2-way ANOVA with Tukey’s multiple comparisons tests. ns = not significant (*p*>0.05). Multiplicity adjusted *p* values are reported.

**Figure S8. Neural mechanisms of negative thermotaxis. a.** CRISPR/Cas9 targeting of the *Ss-tax-4* gene results in iL3s with reduced negative thermotaxis behavior. *Left*: Tracks of no-Cas9 control iL3s and *Ss-tax-4* iL3s migrating for 15 min in a ~17-26°C gradient. T_start_ = 20°C. T_C_ = 23°C. Black dots indicate starting location of each worm. Scale bar = 2 cm, gray zone = 1°C centered at T_start_. A subset of 10 randomly selected tracks are plotted and only a portion of the full gradient is shown. *Right*: Changes in temperature, average speed, and distance ratio exhibited by individual iL3s. n = 26 worms (no-Cas9 control, “Ctrl”), 27 worms (*Ss-tax-4*); *****p*<0.0001, Mann-Whitney test. **b.** HisCl1-mediated silencing of *S. stercoralis* AFD induces deficits in negative thermotaxis. *Left:* Tracks of *Sr-gcy-23.2p*::*strHisCl1*; *Sr-gcy-23.2p*::*GFP* iL3s treated with BU saline or BU saline + 20 or 50 mM histamine for at least 2 h, migrating in a ~22-34°C gradient for 10 min. T_start_ = 20°C. T_C_ = 23°C. Figure conventions are as in panel **a.** *Right*: Change in temperature, average speed, and distance ratio exhibited by individual iL3s. n = 32 worms (BU treated), 33 worms (histamine treated); *****p*<0.0001, Mann-Whitney test. **c.** Diagram of a rapid cooling stimulus spanning the range of temperatures eliciting strong negative thermotaxis in *S. stercoralis* iL3s (22-13°C). Ramp slope: -0.1 °C/s. **d.** *Ce-*AFD and *Ss*-AFD responses to the rapid 22-13°C cooling temperature ramp shown in panel **c**. Colored traces show the responses of individual worms; black traces show median responses. Horizontal dotted lines indicate baseline (R_0_) responses at 22°C. n = 14 worms (*Ce*-AFD), 15 worms (*Ss*-AFD). **e.** Quantification of linear correlation between temperature and calcium responses in a 13-22°C temperature range. Values are Pearson’s correlation coefficients. n = 14 worms (*Ce*-AFD), 15 worms (*Ss*-AFD). ns = not significant (*p*>0.05), Mann-Whitney test. All statistical tests are two-sided.

**Figure S9. Cooling-mediated inhibition in *Ss*-AFD does not display rapid adaptation. a.** *Ss-*AFD calcium responses to a minimum temperature (T_min_) of 13°C following a 22-13°C cooling temperature ramp (Fig. S8c). Colored shading indicates time windows used for analysis. n = 15 worms. T_C_ = 23°C. For calcium responses, colored lines indicate responses from individual worms, black line indicates median response. Temperature traces are mean ± SD recorded temperature (lines + shading; shading may be too small to be visible). Dotted horizontal line indicates baseline temperature response, equivalent to the response at T_C_ (R_Tc_). **b.** *Ss-*AFD steady-state responses at 22°C (following a 15-22°C warming temperature ramp) and 13°C (following a 22-13°C cooling temperature ramp), at early (first 15 s after reaching T_min_ or T_max_) and late (30 s after reaching T_min_ or T_max_ to end of T_min_ or T_max_ stimulus) temporal windows. Responses at 22°C following a 15-22°C warming temperature ramp are replotted from Figure 3. For responses at 22°C, T_C_ = 15°C; for responses at 13°C, T_C_ = 23°C. Despite displaying similar initial magnitudes (early 13°C vs. early 22°C), only the warming-triggered hyperpolarization response at 22°C displays rapid adaptation (early 22°C vs late 22°C); cooling-triggered hyperpolarizations at 13°C are constant across the stated time windows (early 13°C vs late 13°C). ns *=* not significant (*p*>0.05), *****p*<0.0001, 2-way repeated-measures ANOVA with Šidák’s multiple comparisons tests and Bonferroni-Dunn correction; multiplicity adjusted *p* values are reported. Values are median % ΔR/R_Tc_. For responses at 22°C, T_C_ = 15°C; for responses at 13°C, T_C_ = 23°C. Boxes are median and interquartile range; whiskers indicate min and max values. n = 15 worms (13°C), 17 worms (22°C).

**Figure S10. Response of wild-type *C. elegans* ASE neurons to a naturalistic warming temperature ramp.** Responses of *C. elegans* ASER/L neurons expressing YC3.60 to naturalistic warming temperature ramps. Warming stimulus ramp: 20-40°C at 0.025 °C/s. Gray traces indicate responses of individual worms. Black traces show the median response. Temperature traces show the average ± SD recorded temperature (lines + shading; shading is too small to be visible). Horizontal dotted line indicates baseline (R_0_) response at 20°C. T_C_ = 23°C. n = 15 worms. None of the individual traces crossed thermal threshold (T*_ASE_), defined as the first temperature where ΔR/R_0_ deviates from the average *Ce*-ASE_WT_ response by at least 3 * STD of ASE_WT_ ΔR/R_0_ for a minimum of 40 seconds.

**Supplemental Dataset Legends**

The following supplemental data files are available on GitHub:

(<https://github.com/HallemLab/Bryant_et_al_2021>).

**Supplemental Data File 1.** This file includes the results of statistical tests and exact *p* values, as well as the data used for statistical analyses.

**Supplemental Data File 2.** This file includes all primers, plasmids, worm strains, and thermal ramps used in this study.
