## Supplementary material for "The neural basis of temperature-driven host seeking in the human threadworm *Strongyloides stercoralis*": Figure S1

a

| | Ce-GCY-8 | Ce-GCY-18 | Ce-GCY-23 |
|---|---|---|---|
| SSTP_0000354300 (*Ss*-GCY-23.1) | 51.9 | 52.5 | **60.5** |
| SSTP_0000775800 (*Ss*-GCY-23.2) | 49.5 | 51.3 | **57.8** |
| SSTP_0000846800 (*Ss*-GCY-23.3) | 47.2 | 49.0 | **53.3** |

| | Ce-GCY-8 | Ce-GCY-18 | Ce-GCY-23 |
|---|---|---|---|
| SRAE_2000430600 (*Sr*-GCY-23.1) | 50.9 | 52 | **59.6** |
| SRAE_0000007100 (*Sr*-GCY-23.2) | 51 | 51.9 | **59.5** |
| SRAE_X000020900 (*Sr*-GCY-23.3) | 48 | 50 | **54.7** |

b c

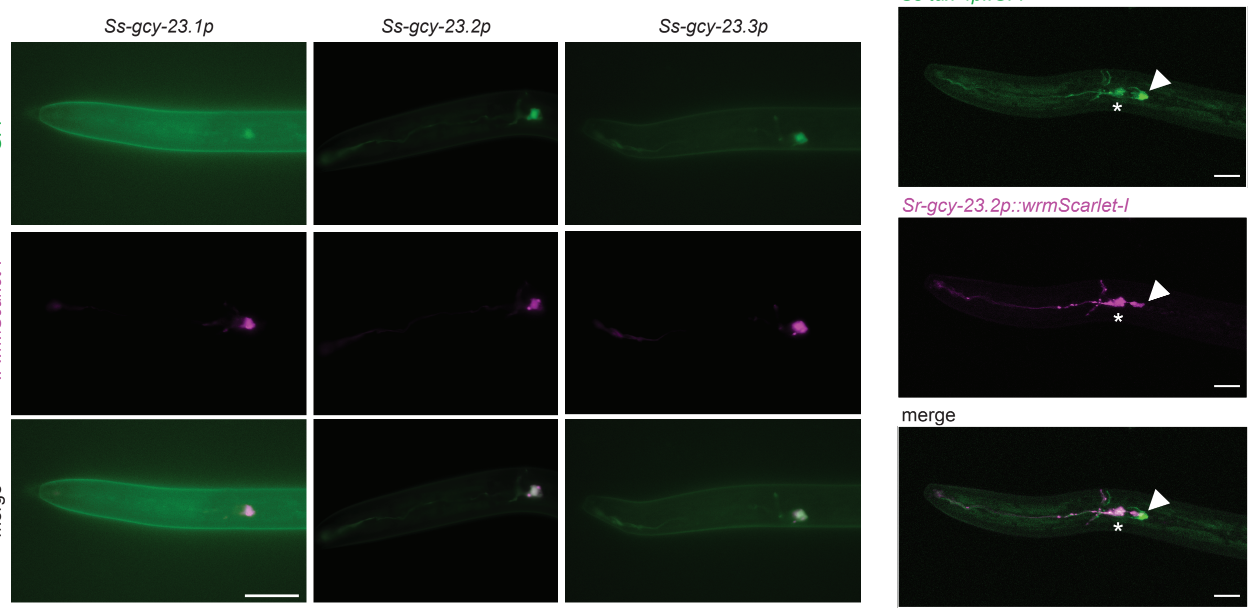
