## Supplementary figures and images for "The neural basis of temperature-driven host seeking in the human threadworm *Strongyloides stercoralis*"

### Figure S2

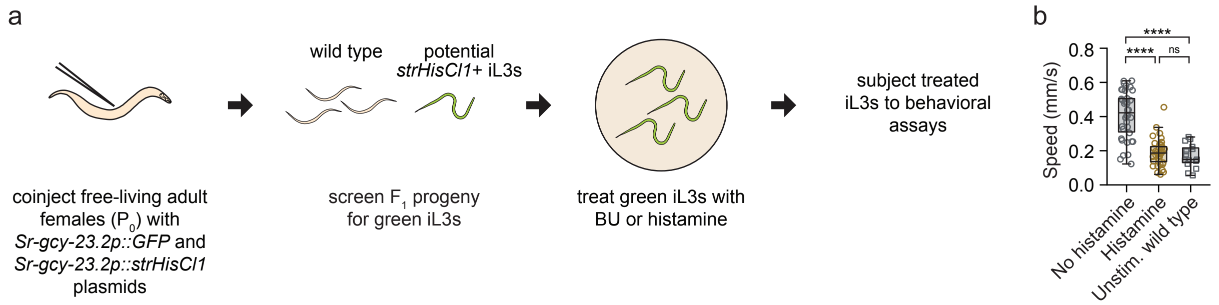

### Figure S3

Figure S3

a

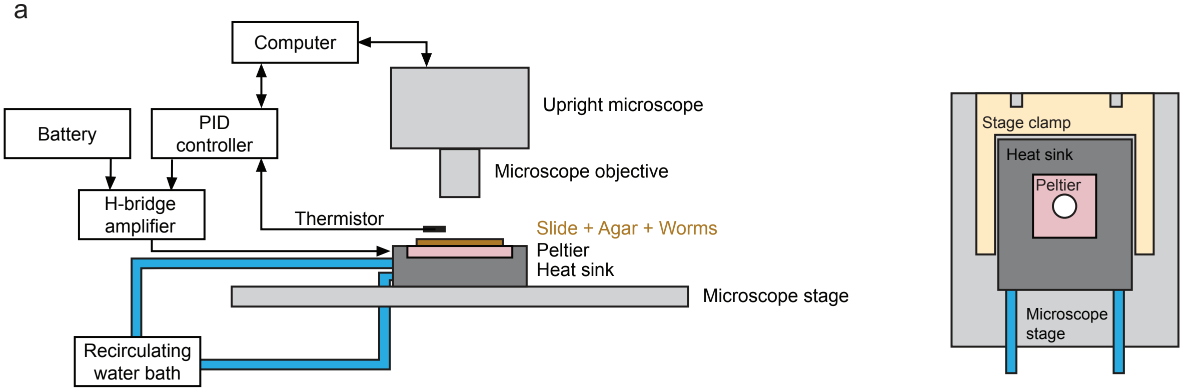


b

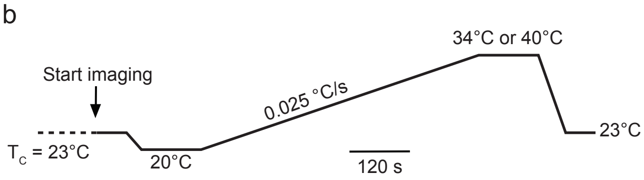


c

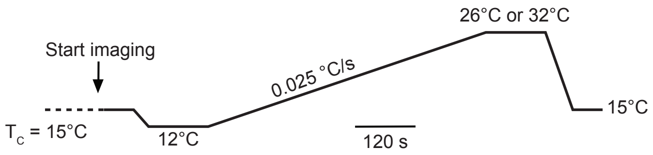

### Figure S4

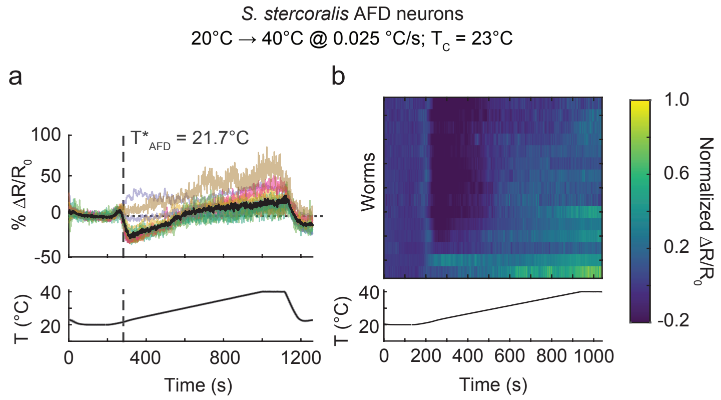

### Figure S5

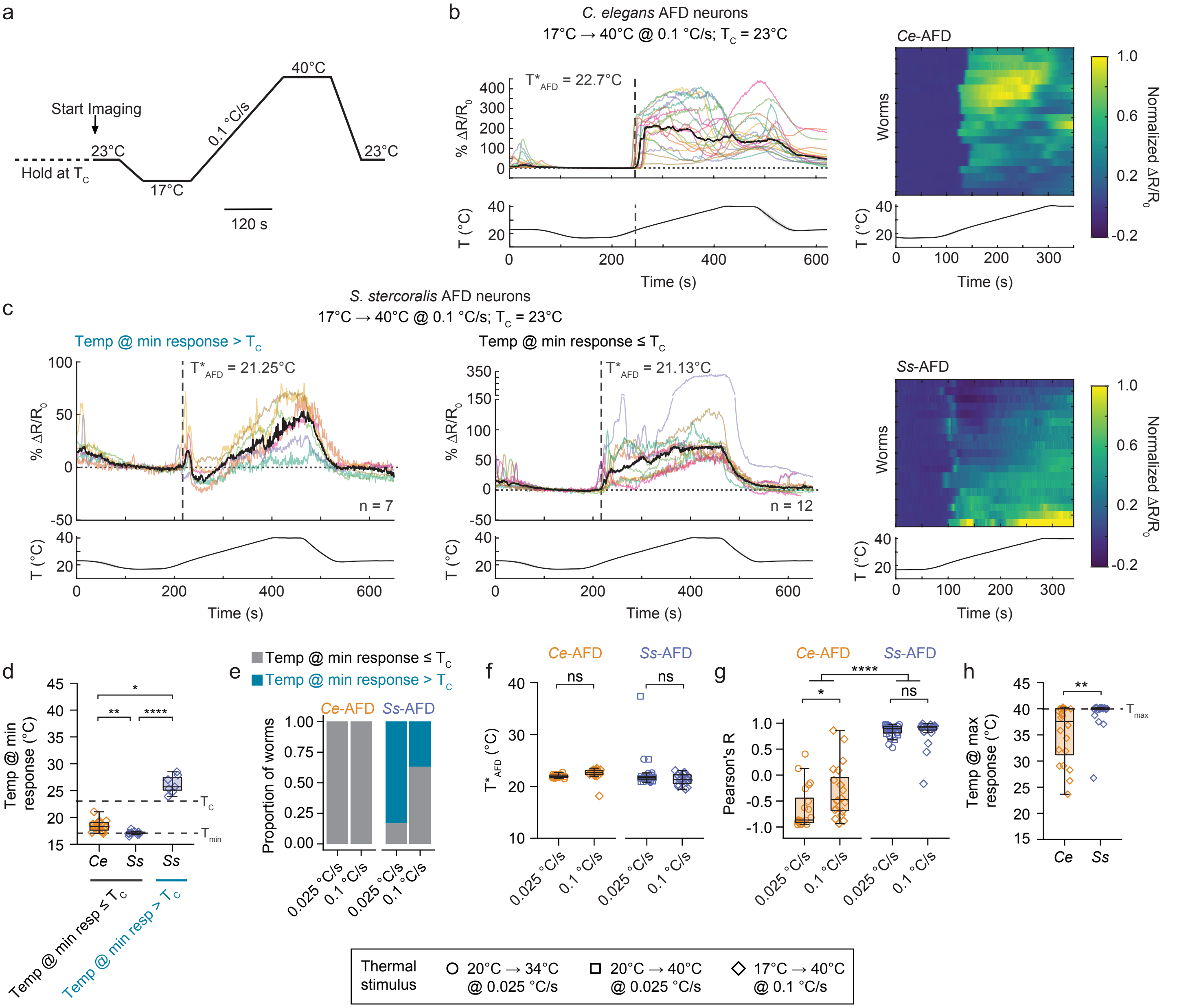

### Figure S6

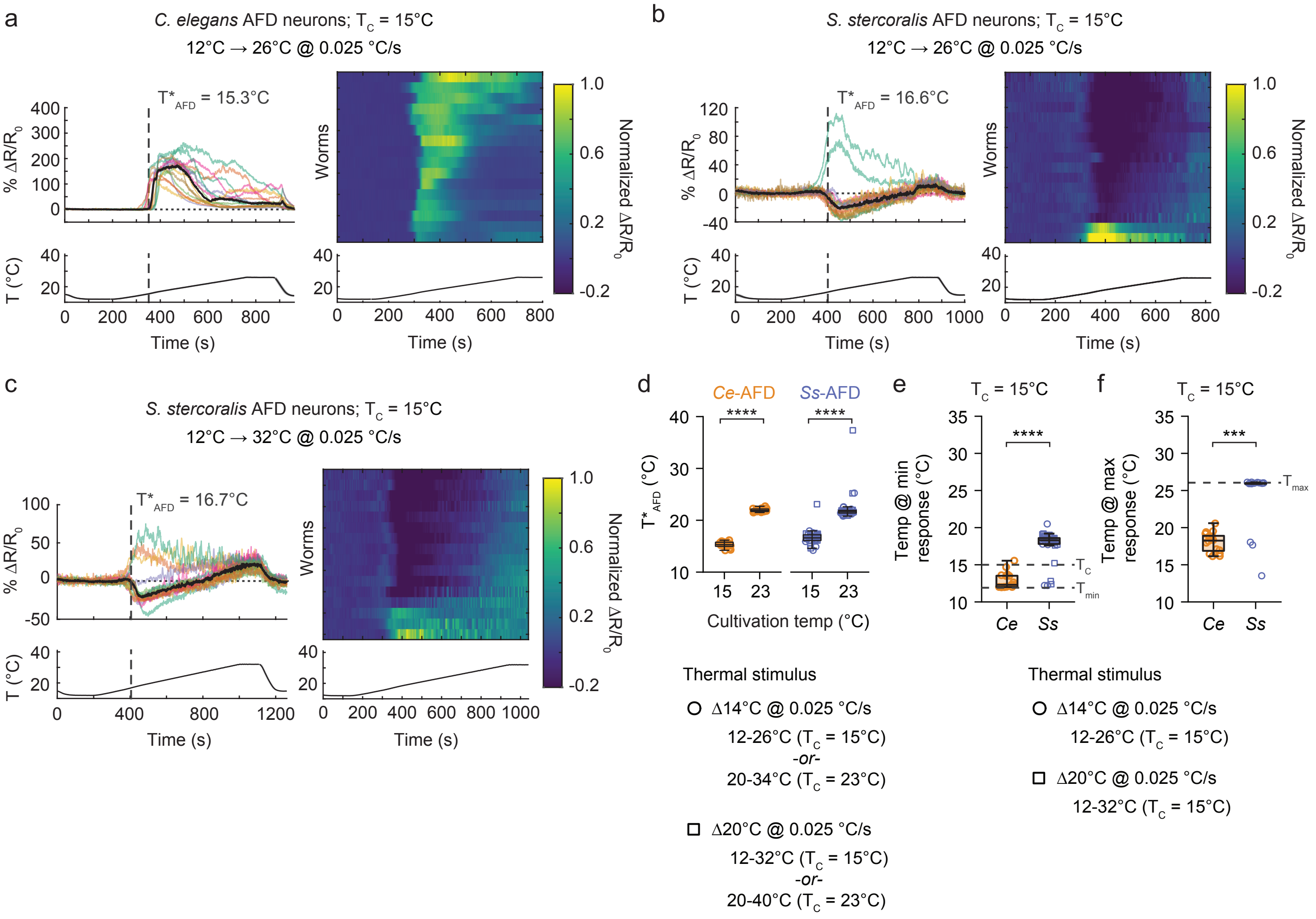

### Figure S7

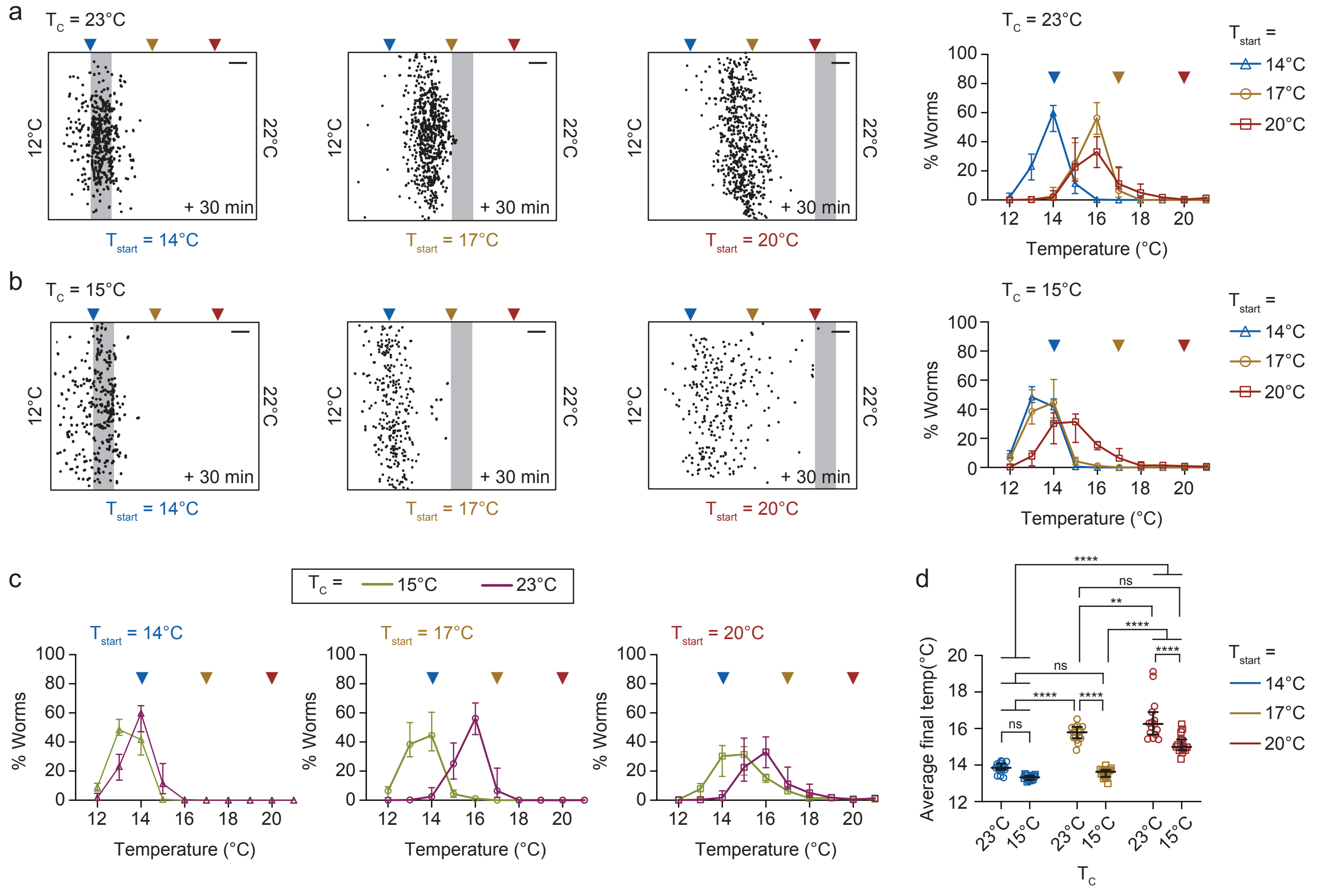

### Figure S8

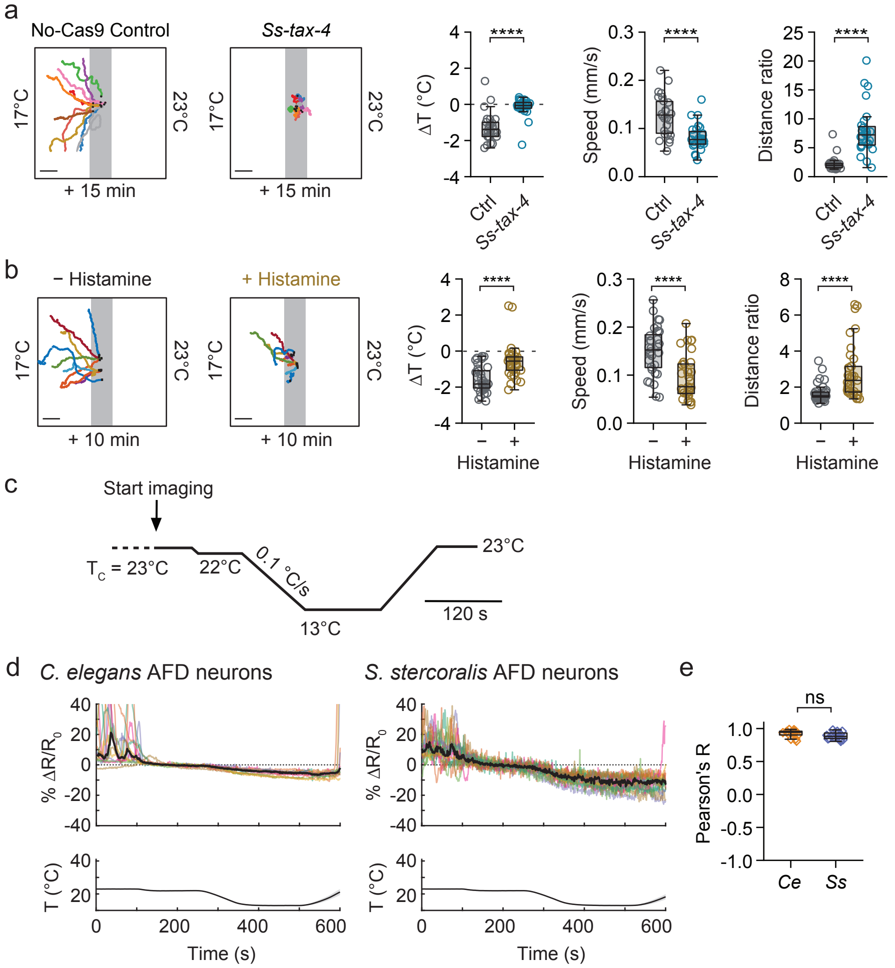

### Figure S9

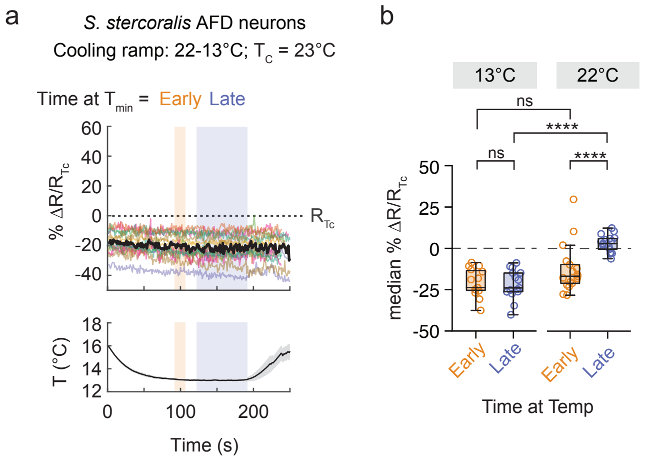

### Figure S10

#### Figure S10

*C. elegans* ASE neurons
20°C → 40°C @ 0.025 °C/s; $T_C$ = 23°C

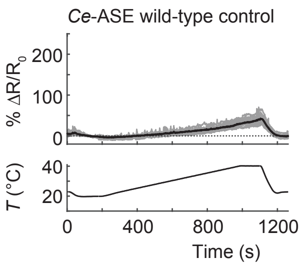
